# From plasmid sequence to process design: A computational analysis of metabolism in the context of plasmid DNA manufacturing

**DOI:** 10.1101/2025.11.12.687962

**Authors:** Nikolaos Stratis, Massimo Morbidelli, Alexandros Kiparissides

## Abstract

Detailed understanding of plasmid-host physiological interactions and recombinant protein expression is crucial for the design and optimization of plasmid DNA (pDNA) manufacturing processes. Successfully transformed host cells carrying one or more copies of a particular plasmid exhibit modified metabolic behavior which may include reduced specific growth rate, increased metabolic resource cycling (ATP/ADP, NAD/NADH, NADP/NADPH) and, in some cases, even modified nutrient uptake and metabolite secretion rates. However, to the extent of our knowledge, there have been no attempts to map the impact of plasmid design directly to critical process parameters (CPPs) such as the specific growth rate and/or productivity. Herein we present, a comprehensive computational analysis based on carbon constrained Flux Balance Analysis (ccFBA) to precisely calculate the metabolic burden imposed by plasmid replication and recombinant protein expression. Explicit stoichiometric coefficients for nucleotides and amino acids were derived directly from the plasmid’s sequence and were used to formulate biosynthetic reactions tailored to the components of individual plasmids. Three tunable parameters were introduced to map the impact of promoter strength, copy number and choice of selection marker on the host cell’s metabolism. Literature derived experimental data from *E. coli* cultures producing three different plasmids were used to constrain the iJR904 Genome Scale Model (GeM). Our analysis revealed correlations between cellular growth rate, promoter strength and pDNA productivity with significant implications for the design of recombinant technology based manufacturing processes.

## 1. Introduction

The increasing number of FDA approved Gene therapies and mRNA based vaccines have created a large demand for plasmid DNA (pDNA) which currently limits global capacity for both classes of therapeutics. Global demand for pharmaceutical grade pDNA is currently projected to reach 10-100 kg by 2030 [1]. However, plasmid propagation and/or expression within the selected microbial host incurs a metabolic burden, due to the additional metabolic resources required for plasmid replication and recombinant protein expression [2],[3],[4],[5]. The magnitude, and therefore impact, of this added resource drain on cellular physiology depends on a number of different factors including plasmid size, copy number, plasmid promoter strength and translation intensity of the derived mRNAs. Elucidation of the resulting process design space via experimentation alone is impractical and time consuming.

Given their wide-spread use in genetic engineering, synthetic biology and recombinant DNA platforms, a huge number of plasmids have been constructed that differ both in terms size and, crucially, sequence. Consequently, significant efforts have been made to study host-plasmid interactions with appropriate computational methods such as Flux Balance Analysis (FBA). FBA has been successfully used to develop custom media formulations tailored to specific plasmids [6], to identify potential pDNA production bottlenecks [4], to simulate protein expression while accounting for plasmid replication and expression intensity [2],[7] and to elucidate potential changes in the metabolic behavior of plasmid bearing *E. coli* cells [8].

Appropriate representation of the stoichiometry of reactions associated with plasmid replication, mRNA expression and translation requires sequence specific manual curation of GeMs. However, the above approaches often include partial (through use of lumped reactions) or generalized stoichiometric representations of plasmid, mRNA, and protein formation. Hatzimanikatis and coworkers [2] augmented the FBA problem formulation through the addition of both proteomic and thermodynamic constraints to develop a high-fidelity computational algorithm that is able to effectively map interactions between the *E*.*coli* host and the plasmid responsible for the additional metabolic burden. Nevertheless, proteomic data is not routinely available and is both time and resource intensive, particularly during early process development.

Herein we present Flux Balance Analysis with host-plasmid interactions (FBAhop), a novel computational algorithm for the automated integration of plasmid-associated reactions into GeMs with stoichiometric coefficients derived directly from the plasmid’s sequence. Literature derived experimental data from E. coli cultures carrying three distinct plasmids were used to constrain the iJR904 GeM which was augmented with sequence specific reactions to account for plasmid replication, mRNA transcription and recombinant protein expression. Design space identification [9] was employed in order to correlate plasmid characteristics to process design. Finally, the metabolic configuration of the resulting plasmid specific GeMs was compared to wild type cells via extensive sampling of the solution space and subsequent analysis with PCA [10] to identify significantly affected pathways. This is the first, to our knowledge, attempt to directly link plasmid sequence to expected process performance paving the way towards truly plug-and-play platform processes.

## 2. Materials and Methods

All simulations were performed using MATLAB 2024b (The Mathworks; Natick, MA, USA), Python (v3.8.18) [11] and the algorithms included in the COBRApy Toolbox [12]. LP optimization problems were solved using the Gurobi solver (version 12.0) [13].

### 2.1 Carbon constrained flux balance analysis (ccFBA)

FBA, a well established constraint-based modelling approach, calculates the distribution of metabolic resources (carbon, energy, redox) across all metabolic reactions included in a GeM. Biochemical reactions are represented in the form of a stoichiometric matrix ***S*** of size (m x n). Every row (*m*) of ***S*** represents a unique compound, while every column (*n*) represents a single reaction. Consequently, each element (***sij***) of ***S*** contains the stoichiometric coefficient of the *i*^*th*^ metabolite in the *j*^*th*^ reaction. The flux through all (*n*) reactions is represented by the (n x 1) vector ***v***. Assuming that cells reach a pseudo-steady state, where the net concentration change (production minus consumption) for each metabolite in the network is equal to zero, a mass balance across the entire metabolic network yields:

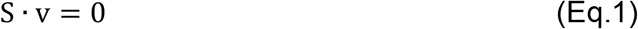

Equation (1) defines an underdetermined system of algebraic steady-state mass balance equations, as the number of unknown reaction fluxes (***v***), exceeds the number of available balance equations (m). In order to be able to calculate all the individual reaction fluxes in (***v***), FBA-based approaches reformulate the problem as a linear programming (LP) optimization problem under the assumption that cells configure their metabolism in a manner that seeks to optimize a particular objective [14]. Therefore, the problem can be summarized by Equations (2-4):

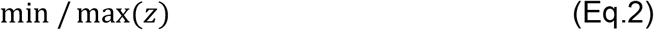

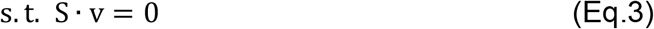

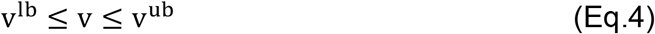

where z corresponds to the flux or sum of fluxes being optimized subject to mass balance constraints (Equation 3) and the set of inequality constraints (Equation 4). In the present study, maximization of biomass was used as the objective function (z = ν_biomass_). The set of inequality constraints described by Equation 4 is used to constrain flux values between an upper (ν^ub^) and lower (ν^lb^) bound. These reaction bounds can be significantly refined by constraining the permissible flux through intracellular reactions based on the amount of carbon taken up by the cell under the studied physiological conditions [15]. Therefore, under carbon constrained FBA (ccFBA), Equation (4) is replaced by:

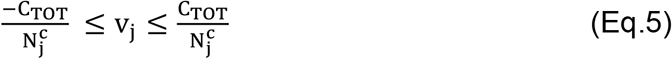

where *C*_*TOT*_ is the total carbon flux flowing into the cell and 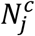 is the number of the carbon atoms that participate in reaction j.

### 2.2 GeM selection and tuning

Several GeMs for *E. coli* have been published in the scientific literature, including highly curated community models. The latest version of four *E. coli* GeMs was obtained from the BiGG database [16] (Table 1) and the ability of each model to accurately predict intracellular fluxes was assessed against experimental data.

**Table 1.** Information of Genome Scale Models benchmarked in this study.

| GeM | Metabolites | Reactions | Genes | Reference |
| --- | --- | --- | --- | --- |
| iJR904 | 761 | 1075 | 904 | [17] |
| iAF1260 | 1109 | 1285 | 987 | [18] |
| iJO1366 | 1805 | 2583 | 1367 | [19] |
| iML1515 | 1877 | 2712 | 1516 | [20] |

^13^C metabolic flux data for intracellular reactions across several central carbon pathways including Glycolysis, Pentose Phosphate Pathway, Entner-Doudoroff Pathway, TCA Cycle, Glyoxylate Shunt, Amphibolic Reactions and Amino Acid Biosynthesis were retrieved from literature [21]. The dataset contained experimentally measured values for intra- and extra-cellular reaction fluxes across different environmental conditions (aerobic and anaerobic) and substrates (Glucose and Xylose). Each GeM was constrained using experimental values for uptake and secretion rates, while the specific growth rate was predicted using ccFBA. The resulting solution space was uniformly sampled with an Artificial Centering Hit and Run (ACHR) algorithm [22]. Mean flux values were derived from the sampled fluxes and compared against experimental data. The mean absolute error across all reactions was calculated and normalized to the largest error obtained among all models, resulting in error values scaled between 0 and 1. This process was repeated once for each combination of GeM, environmental condition, and substrate, resulting in 16 distinct combinations. Additionally, each model was benchmarked based on its ability to predict the correct directionality of the measured reaction fluxes. Models that failed to predict the correct direction were assigned a penalty point.

The iJR904 GeM displayed the lowest normalized absolute error across each of the four conditions examined (Figure 1). In addition, the total number of reactions across all conditions for which a GeM failed to correctly predict reaction directionality was tracked, with iJR904 accumulating the lowest number of mismatches (Supplementary Figure S1).

**Figure 1.**
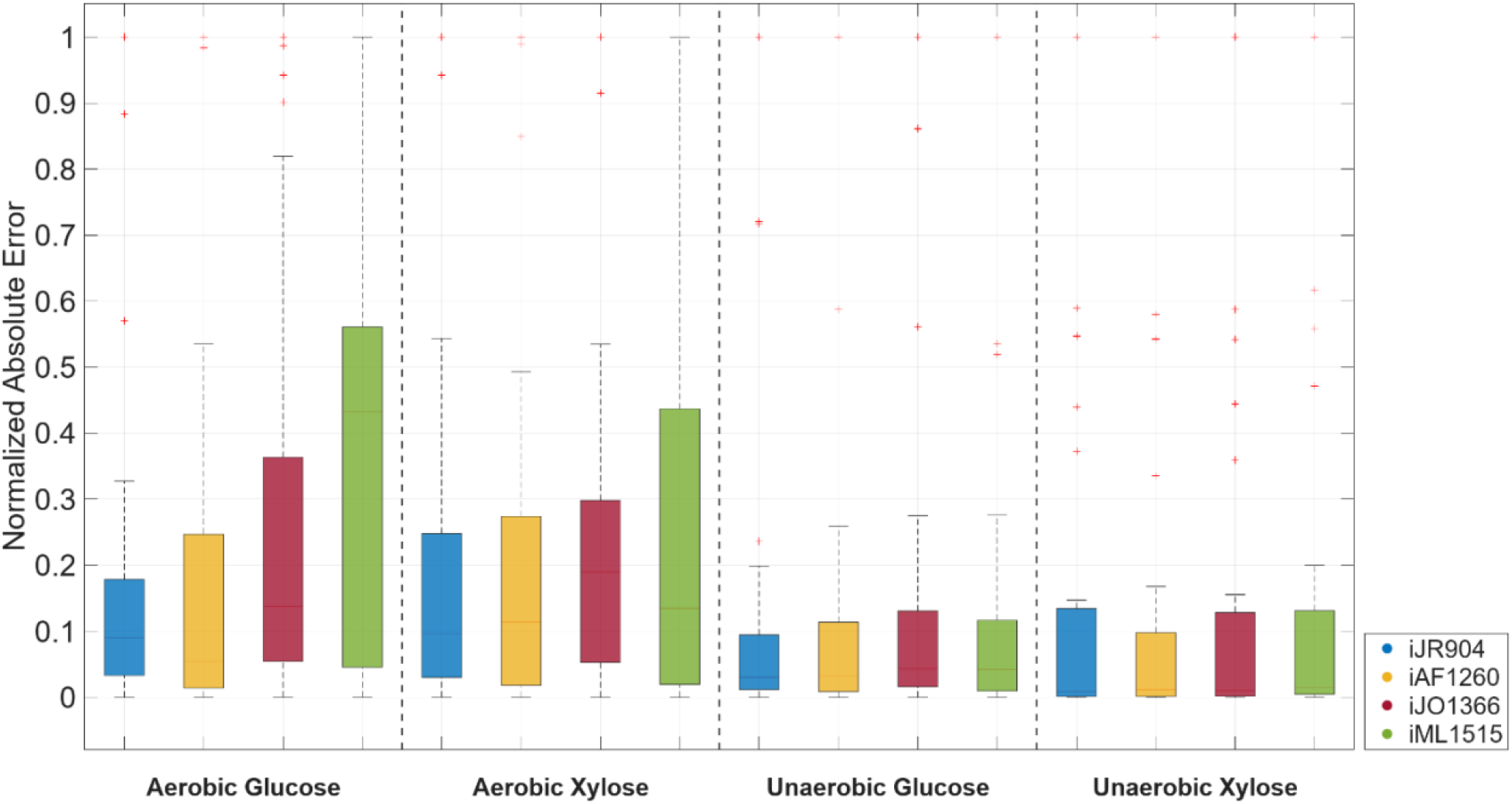
Normalized Absolute Error between experimentally measured fluxes from [21] and the distribution means of sampled flux values

### 2.3 Flux balance analysis with host plasmid interactions (FBAhop)

Initially, GenBank files in FASTA format for each plasmid are parsed to extract and enumerate exact nucleotide sequences. Briefly, open reading frames (ORFs) with a minimum length of 75 nucleotides are identified using the orffinder tool [23], while nested ORFs are removed. The nucleotide sequences of identified ORFs are extracted, transcribed into mRNA sequences and translated into the corresponding protein sequences. Subsequently, translated ORFs are subjected to BLASTP using the NCBIWWW.qblast tool [23], and the resulting encoded protein names are obtained and displayed. As a result the exact number and composition of DNA and RNA nucleobases as well as the number and composition of required amino acids for plasmid replication and expression is computed.

Therefore, a unique, plasmid specific replication reaction with tailored stoichiometry based on the parsed plasmid sequence can be developed. The general structure of the plasmid replication reaction can be summarized by Equation 6:

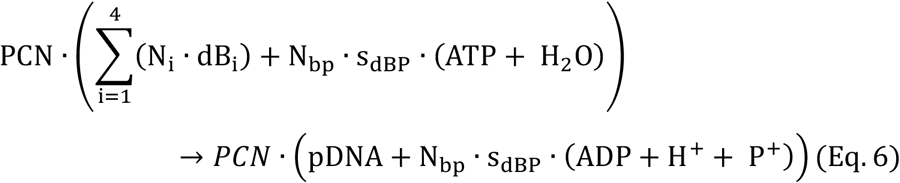

where (*PCN*) is the plasmid copy number, (N_i_) is the total number of the *i*^*th*^ DNA base (dB_i_) included in the plasmid’s sequence, (N_bp_) is the total number of base pairs and (s_dBP_) is the average ATP and H_2_O cost of synthesis per mol of nucleotide base.

Accordingly the mRNA transcription reaction can be summarized by Equation 7:

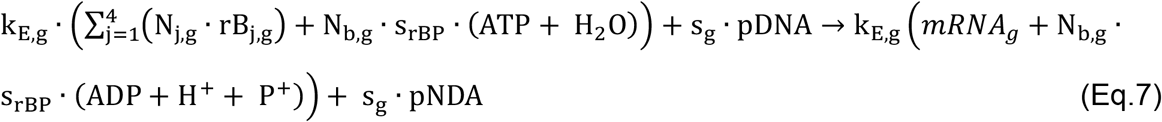

where (k_E,g_) is a parameter representing the strength of the promoter under which gene (*g*) is expressed 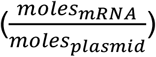, (N _j,g_) is the total number of the *j*^*th*^ RNA base (rB_j_) translated from gene *g’s* sequence, (N_b,g_) is the size of gene *g* (bases), (s_rBP_) is the average ATP and H_2_O cost of synthesis per mol of ribonucleotide base and (s_g_) is a binary variable equal to 1 if gene (*g*) encodes a selection pressure marker (such as an antibiotic resistance protein) and equal to 0 otherwise. Previous studies have shown that a higher flux toward the expression of an antimicrobial resistance protein can lead to increased pDNA production [24], however no such correlation has been established for the expression of recombinant proteins not tied to the cell’s survival.

Finally, protein synthesis is summarized by Equation 8

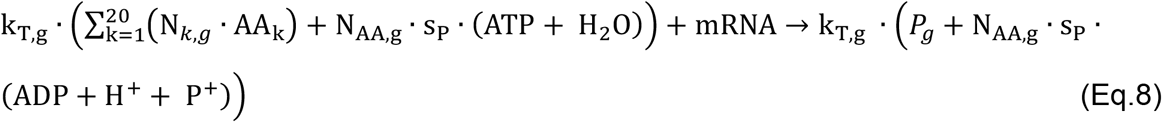

where (k_T,g_) is a parameter representing the translation efficiency of mRNA transcribed from gene (*g*) to protein 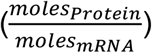, (N_k,g_) is the total number of the *k*^*th*^ amino acid (AA_k_) contained in the protein’s sequence, (N_AA,g_) is the total number of amino acids contained in the recombinant protein encoded by gene (*g*) and (s_P_) is the average ATP and H_2_O cost of synthesis per mol of amino acid. mRNA is treated solely as a reactant in Equation 8 due to its short half-life (2–10 minutes) [25].

The ATP hydrolysis cost in equations 6-8 is determined by multiplying the number of molecular building blocks (N_bp_, N_b,g_ and N_AA,g_ respectively) by the ATP requirement per mole of building block, as described in [26] and summarized in Table 2.

**Table 2.**
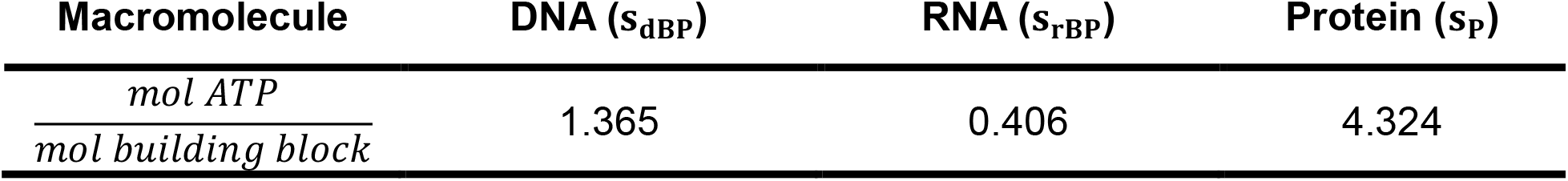
Energetic requirements for macromolecule biosynthesis for DNA, RNA and Protein.

#### Integration of plasmid associated reactions into GeMs

The reaction described by equation 6 and as many copies of the reactions described by equations 7-8 as there are genes on the plasmid are added to the host cell organism’s GeM. To ensure connectivity with the metabolic network and consistent mass balancing, demand reactions are also added for pDNA and expressed recombinant proteins (*P*_*g*_) not associated with a selection marker. Recombinant proteins associated with a selection marker, such as antibiotic resistance proteins, are instead directly incorporated into the GeM’s biomass equation since they are necessary for the cell’s survival [27]. Nevertheless, users of the FBAhop toolbox maintain the ability to choose whether a particular recombinant protein should be incorporated into the biomass equation or not.

When a recombinant protein is incorporated directly into the biomass equation, the stoichiometric coefficients for all amino acids in the biomass equation are adjusted to account for the relative abundance of amino acids in the recombinant protein according to equation 9.

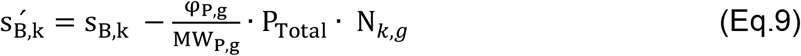

where: (s_B,k_) is the stoichiometric coefficient of the k^th^ amino acid (AA_k_) in the biomass equation, (φ_P,g_) is the heterologous protein’s weight fraction 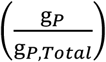 in respect to the total protein content of the cell (P_Total_), (MW_P,g_) is the heterologous protein’s molecular weight and (N_k,g_) is the total number of the *k*^*th*^ amino acid (AA_k_) contained in the protein’s sequence. It is important to note that φ_P,g_ is not directly derived from the protein flux calculated in Eq.8 as a portion of the translated protein may form aggregates, misfold, or be secreted [28].

The FBAhop algorithm is freely available as a COBRApy compatible toolbox at https://github.com/alex-kip/FBAhop. FBAhop dependencies include COBRApy, the Biopython package and the orfinder tool.

## 3. Results & Discussion

The developed algorithm was employed to investigate the impact of three different plasmids on *E. coli* metabolism and physiology. Specifically, uptake and secretion rates from batch *E. coli* cultures transformed with the pOri1 (2811 bp), pOri2 (4575 bp) [29] and pMal-p2x (6720 bp) [7] plasmids were obtained from literature. All three plasmids are equipped with an ampicillin resistance gene which encodes the β-lactamase protein. In addition, the pMal-p2x plasmid carries the lacI gene that encodes the lac repressor protein under a lacI^q^ promoter. The lac repressor protein reversibly binds a lac operator immediately downstream of a P_tac_ promoter thus regulating the inducible expression of a Glucose Isormerase-Maltose Binding Protein fusion protein (GI-MBP). The above set of plasmids allowed us to study both plasmid replication (pOri1, pOri2) in the context of pDNA manufacturing and plasmid replication coupled with recombinant protein expression (pMal-p2x).

### 3.1 Parameter Tuning and Analysis

Initially, the iJR904 GeM was constrained with literature derived experimental data from *BL21 E. coli* cultures carrying each of the three plasmids (Table 3), thus generating three distinct models. The three plasmid sequences were parsed and stoichiometrically tailored equations for plasmid replication (Eq. 6; pOri1, pOri2, pMal-p2x), antibiotic resistance protein expression (Eq. 7-8; β-lactamase) and recombinant mRNA/protein expression (Eq. 7-8; lacI, GI-MBP) were developed and added to the respective models. Each model was further refined via the estimation of carbon-flux based constraints for intracellular reactions using ccFBA [15].

**Table 3.** Literature derived uptake and secretion rates used in the present study.

| Exchange rates<br>(mmol·gDW<sup>-1</sup>·h<sup>-1</sup>) | pOri1 |  | pOri2 |  | pMal-p2x |  |
| --- | --- | --- | --- | --- | --- | --- |
|  | LB | UB | LB | UB | LB | UB |
| **PCN** | 50 | 50 | 410 | 410 | 10 | 10 |
| **Glucose** | -7.35 | -6.65 | -6.61 | -5.98 | -1.26 | -1.14 |
| **Acetate** | 3.89 | 4.30 | 4.18 | 4.62 | 0.38 | 0.42 |
| **Oxygen** | -17.74 | -16.05 | -17.32 | -15.67 | -3.46 | -3.13 |
| **CO<sub>2</sub>** | 17.00 | 18.79 | 16.43 | 18.16 | - | - |
| **lacl** | - | - | - | - | 4.35e-6 | 1.30e-5 |
| **GI-MBP** | - | - | - | - | 8.26e-6 | 9.13e-6 |
| **Reference** | [29] |  | [29] |  | [7] |  |

Both the pOri1 and pOri2 plasmids do not express any recombinant protein products apart from the antibiotic resistance protein, β-lactamase. As mentioned in section 2.3 (Eq. 8-9) the antibiotic resistance protein is incorporated into the GeM’s biomass reaction which is readjusted according to the weight fraction of the resistance protein (*φ*_*P*_). Therefore the final problem formulation for the pOri1 and pOri2 models is described by equations Eq. 10-14:

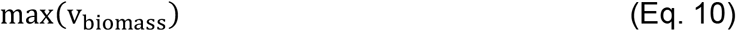

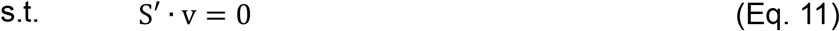

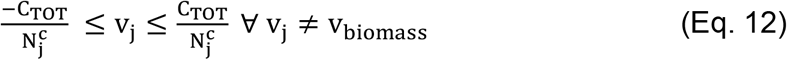

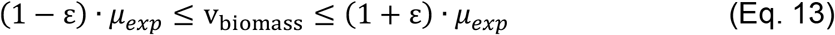

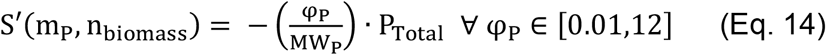

where S’ is the iJR904 stoichiometric matrix augmented by the addition of the reactions for plasmid replication (Eq. 6), antibiotic resistance gene expression (Eq. 7) and translation (Eq. 8) as well as a demand reaction for the produced pDNA. Upper and lower bounds for intracellular reactions are computed via ccFBA (Eq. 12), while the biomass reaction was bound to the experimentally reported value (*μ*_*exp*_) with a small tolerance margin (*ε*, Eq. 13). Finally, the biomass equation of the iJR904 GeM is adjusted via the addition of the antibiotic resistance protein (P) as a reactant (Eq. 14) and the stoichiometric coefficients for all amino acids in the biomass equation (s_B,k_) are adjusted according to Eq. 9.

Direct experimental measurement of expression (*k*_*E*_) and translation efficiencies (*k*_*T*_) is both resource intensive and not practical considering the large number of available plasmids. Instead, a global parametric analysis was performed in order to compare each model’s output when simulated with randomly sampled parameter values against the experimental data of Table 3. A feasible range for each FBAhop parameter (*k*_*E*_, *k*_*T*_, *φ*_*P*_) was defined based on relevant scientific literature. Specifically, *k*_*E*_ was set between 0.02 and 3 mRNA molecules per gene 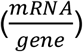 [30], *k*_*T*_ was set between 10 and 100 protein molecules per mRNA molecule 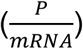 [31] and *φ*_*P*_ was set between 0 and 12% total protein content 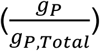 [32]. The parameters were then allowed to vary simultaneously within the above defined ranges. A total of 2^20^ points were sampled from the resulting multi-dimensional parameter space using a Sobol sequence based random number generator [33]. Each of the three models (pOri1, pOri2, pMal-p2x) was simulated once for each set of sampled parameter values and parameter sets that accurately predicted the experimentally reported growth rate were recorded.

The expression efficiency (*k*_*E*_) values that yield predicted growth rates in agreement with experimental data for both the pOri1 and the pOri2 plasmids form a well defined skewed normal-like unimodal distribution (Figures 2A, 3A). In contrast the distributions of selected values both for translation efficiency (*k*_*T*_) and for the fraction of total protein that corresponds to the antibiotic resistance protein (*φ*_*P*_) form exponential-like distributions for both the pOri1 and pOri2 plasmids (Figures 2B-C, 3B-C). In fact values towards the lower bound set for translation efficiency (*k*_*T*_) have a higher probability of accurately predicting the experimental growth rate (Figures 2B, 3B), while the inverse is true for the recombinant protein fraction (*φ*_*P*_, Figures 2C, 3C).

**Figure 2.**
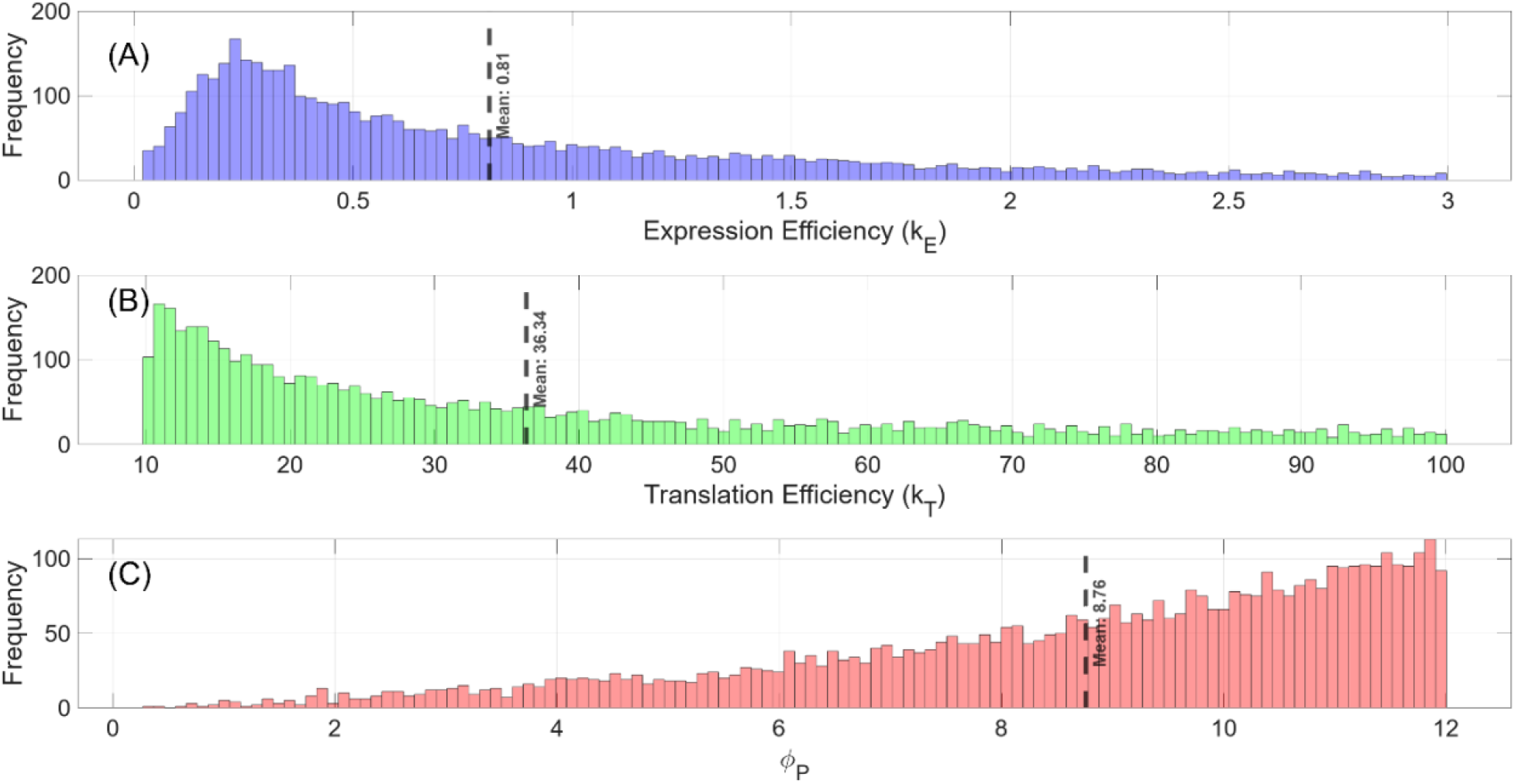
Global parametric analysis of FBAhop parameters for plasmid pOri1. Distributions of randomly sampled values that can accurately predict experimentally reported growth rates: A) expression efficiency (k_E_ Eq.7); B) translation efficiency (k_T_ Eq.8) and fraction of total protein corresponding to the antibiotic resistance protein (φ_P_)

**Figure 3.**
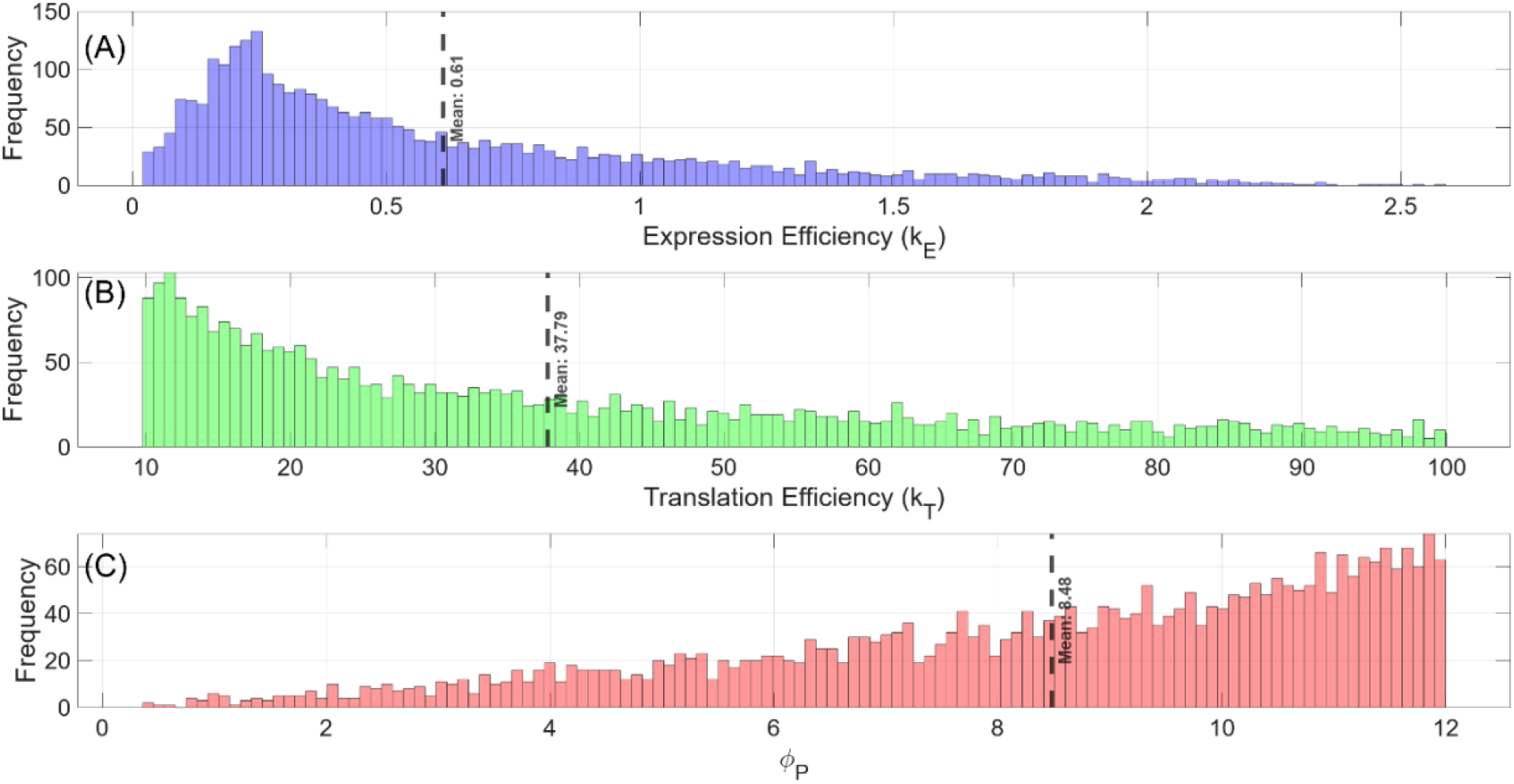
Global parametric analysis of FBAhop parameters for plasmid pOri2. Distributions of randomly sampled values that can accurately predict experimentally reported growth rates: A) expression efficiency (k_E_ Eq.7); B) translation efficiency (k_T_ Eq.8) and fraction of total protein corresponding to the antibiotic resistance protein (φ_P_)

Interestingly, in all cases distribution means align well with experimentally reported values for endogenous gene expression (Figures 2, 3). Specifically, the mean expression efficiency (*k*_*E*_) value for pOri1 is 0.81 mRNA molecules per plasmid and 0.61 for pOri2 which compare well with experimentally observed values for bacterial cells (0.4 transcripts per gene per cell [30]). Similarly, the average recombinant protein fraction (*φ*_*P*_) is consistent with experimentally observed values 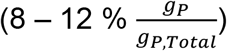 [32].

For the pMal-p2x plasmid, FBAhop parameterization was extended to include three distinct gene modules: the antibiotic resistance gene encoding β-lactamase, the lac repressor regulatory gene (lacI), and the recombinant fusion protein product genes (GI-MBP). Each gene module was assigned unique transcriptional (*k*_*E*_) and translational (*k*_*T*_) efficiency parameters to accurately capture their differential expression profiles. To ensure flux through the recombinant protein reaction a simple weighted multi-objective optimization problem was formulated. Both terms in the objective function were normalized by their corresponding experimental reference values, enabling an equal assessment of the trade-off between biomass growth and recombinant protein production. Consequently, the multi-objective optimization problem was formulated to simultaneously maximize the scaled biomass and recombinant protein fluxes:

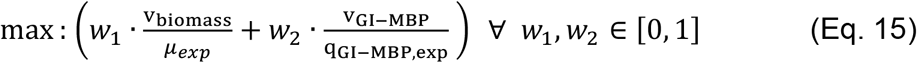

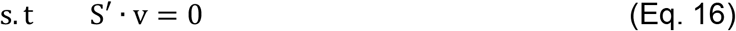

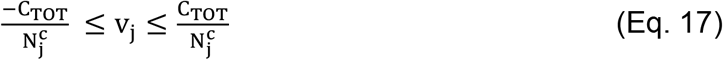

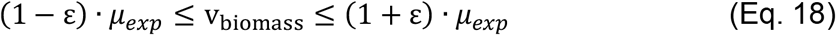

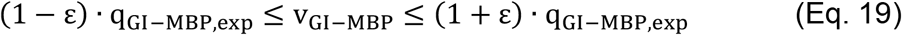

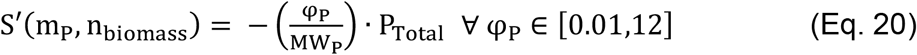

where S’ is the iJR904 stoichiometric matrix augmented by the addition of the reactions for plasmid replication (Eq. 6), antibiotic resistance gene expression (Eq. 7) and translation (Eq. 8). Eq.6-8 were added once for each of the three expressed genes on the pMal-p2x plasmid, namely the genes encoding β-lactamase, lacI and GI-MBP. Demand reactions for the produced pDNA,lacI repressor and recombinant fusion protein product (GI-MBP) were also added to S’. Upper and lower bounds for intracellular reactions are computed via ccFBA (Eq. 17), while the biomass and recombinant fusion protein reactions were bound to the experimentally reported values (*μ*_*exp*_ and q_GI™MBP,exp_ respectively) with a small tolerance margin (*ε*, Eq. 18-19). Finally, the biomass equation of the iJR904 GeM was adjusted via the addition of the antibiotic resistance protein (P) as a reactant (Eq. 20) and the stoichiometric coefficients for all amino acids in the biomass equation (s_B,k_) were adjusted according to Eq. 9.

Interestingly, different behavior was observed in simulations carrying the more complex pMal-p2x plasmid. While distribution means for parameters (*k*_*E*_, *k*_*T*_, *φ*_*P*_) related to the expression of the antibiotic resistance marker were comparable to those found in pOri1 and pOri2 simulations, distribution means for parameters (*k*_*E*_, *k*_*T*_) related to the recombinant products (lacI and GI-MBP) were significantly higher (Supplemental Figures S2, S3). In addition, parameter values related to the recombinant products, that could successfully predict the experimentally reported growth and GI-MPB protein secretion rates appeared to be uniformly distributed and did not follow any discernible pattern. This variability likely results from the added complexity of the pMal-p2x plasmid, which encodes multiple functionally distinct gene modules (the regulatory lac repressor, the antibiotic resistance gene and fusion protein product) with independent and potentially interacting transcriptional and translational controls which are not represented in the present model formulation. Furthermore, the multi-objective optimization formulation (Eq. 15) incorporating both biomass growth and recombinant protein flux creates additional trade-offs that broaden feasible parameter distributions. Such complexity is typical in biological systems with layered regulation and varied expression modalities. Consequently, parameter distributions for pMal-p2x span a wider space without strong convergence, reflecting inherent biological and modeling uncertainties.

### 3.2 Metabolic Fingerprint of plasmid expression and replication

To further understand the impact and functional role of the parameters introduced by FBAhop a series of *in silico* experiments was performed. Specifically, a 3-level Box Behnken Design was employed to generate sets of parameter values for expression (*k*_*E*_), translation (*k*_*T*_) and recombinant protein weight fraction (*φ*_*P*_) parameters. The levels for the experimental design were based on the sampled distribution means ± one standard deviation (Figures 2, 3 and Supplemental Figures S2, S3). Each of the parameter sets defined by the experimental design (Supplemental Tables S1, S2) were simulated separately. The resulting models were carbon constrained [15] and 10,000 solution vectors were retrieved from each model’s solution space using a Markov-Chain Monte Carlo algorithm [22]. This resulted in 15 unique models simulating *E. coli* cells carrying the pOri1 plasmid, 15 unique models for cells carrying the pOri2 plasmid and 62 unique models for cells carrying the pMal-p2x plasmid.

Principal Component Analysis (PCA) was employed to systematically assess the metabolic reconfiguration of host cells following plasmid transformation and recombinant product expression following the methodology presented by [10]. Briefly, the sampled solution vectors from the models generated by the experimental design were allocated in data pools according to the carried plasmid. Each data pool was augmented by the addition of solution vectors from wild type *E. coli* carrying no plasmid. PCA was applied individually to each data pool and loadings with a value higher than 90% of the highest loading were recorded for further analysis.

PCA revealed that plasmid bearing cells display distinct metabolic characteristics compared to wild type *E. coli* cells since the respective sampled solution vectors are distinctly separated, with most of the variance captured by the first two principal components (Figures 4, 5). This clear separation suggests multiple reaction fluxes, beyond those associated with plasmid replication and expression differentiate the metabolic configuration of transformed cells from wild-type metabolism. The first principal component (PC) alone contains enough information to distinguish wild type from plasmid bearing cells, while the first four PCs can clearly distinguish between cells carrying plasmids with different translation and expression characteristics.

**Figure 4.**
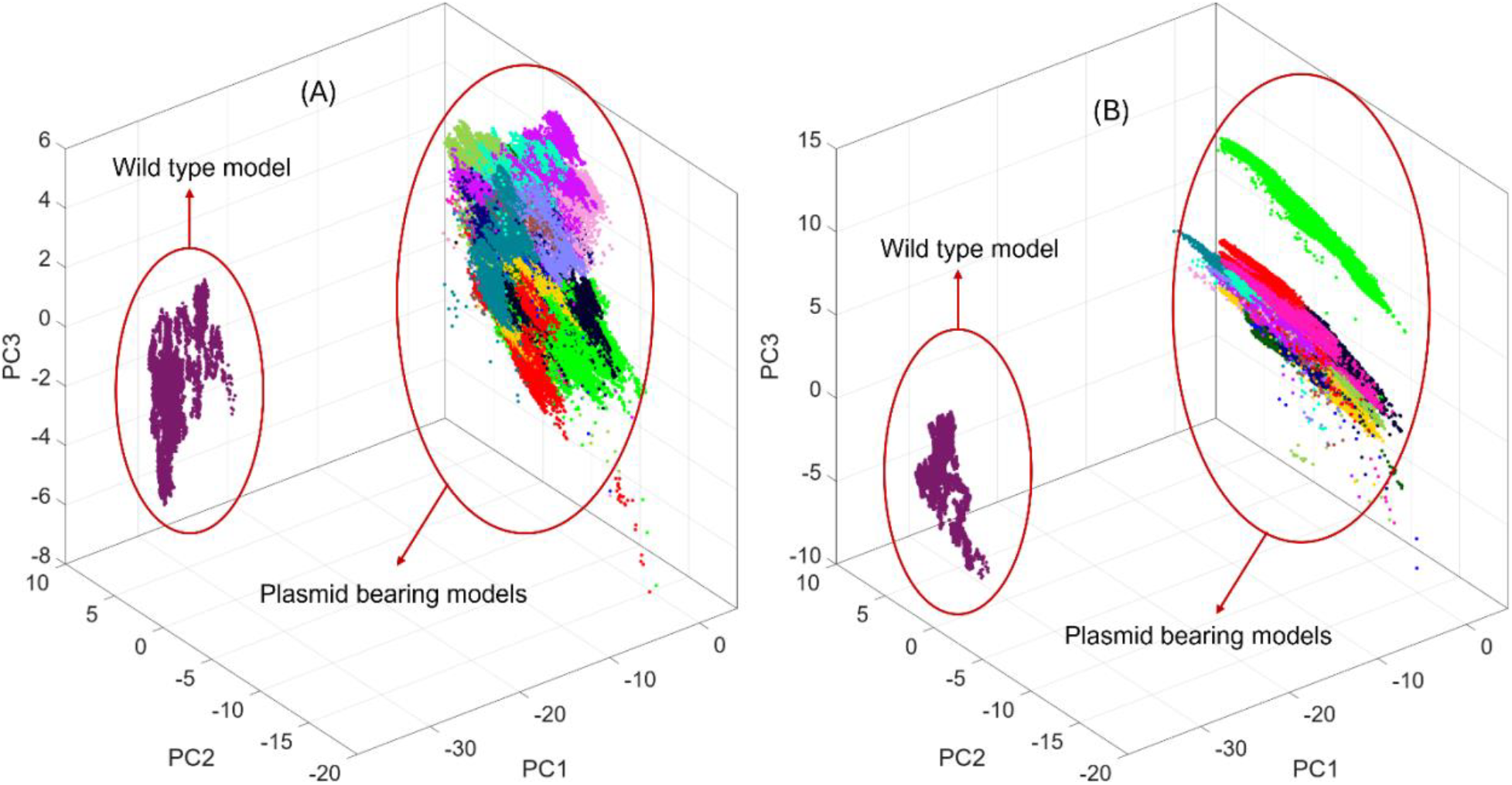
Metabolic fingerprint of plasmid bearing *E. coli* cells. Principal component analysis of sampled solution vectors retrieved from both wild type and plasmid bearing simulations with varying parameter values for the FBAhop parameters (*k*_*E*_, *k*_*T*_, and *φ*_*P*_) for cells carying the A) pOri1 plasmid and B) the pOri2 plasmid.

**Figure 5.**
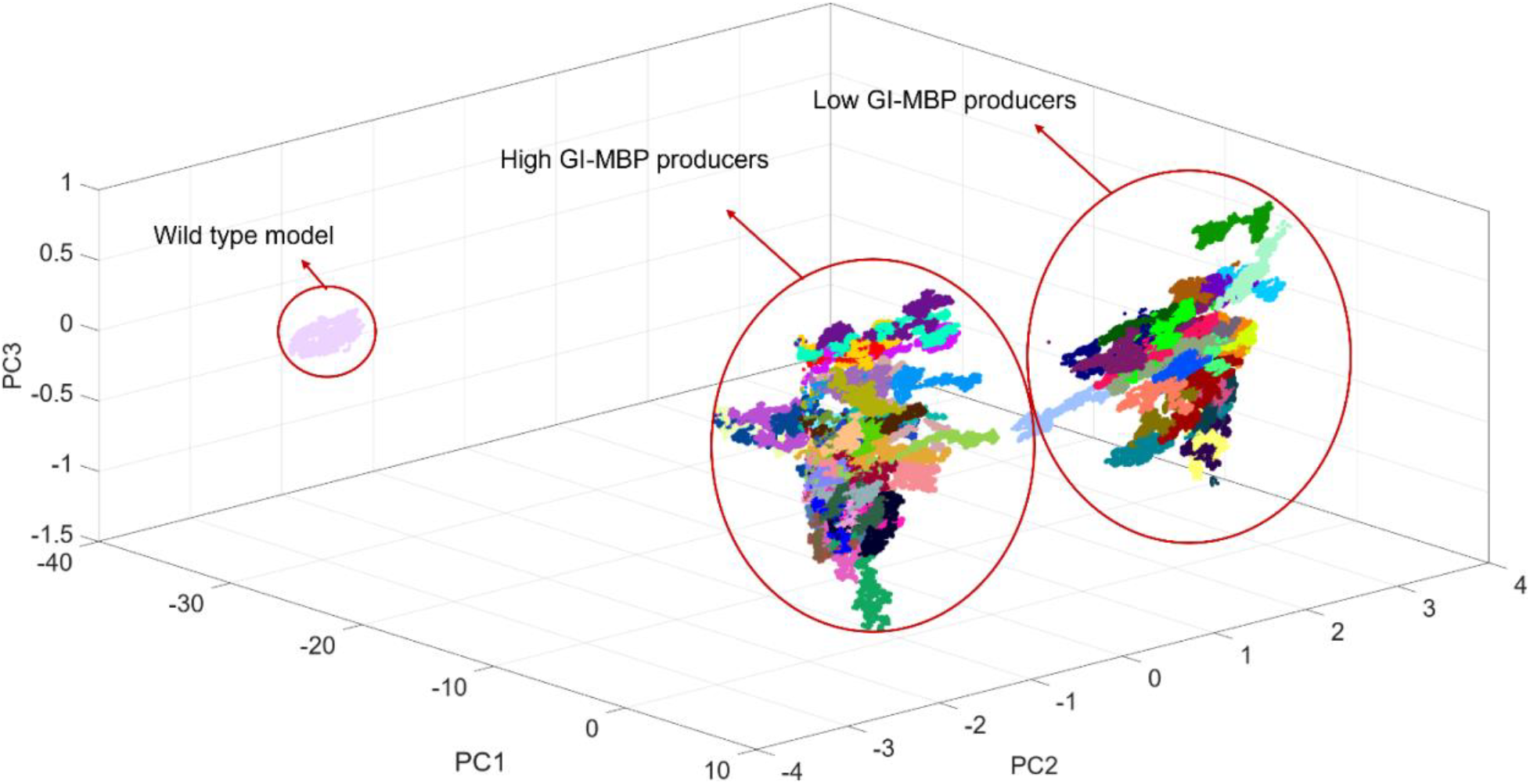
Metabolic fingerprint of plasmid bearing *E. coli* cells. Principal component analysis of sampled solution vectors retrieved from both wild type and plasmid bearing simulations with varying parameter values for the FBAhop parameters (*k*_*E*_, *k*_*T*_, and *φ*_*P*_) for cells carying the pMal-p2x plasmid.

The first 31 PCs for simulations carying the pOri1 and pOri2 plasmids and the first 8 PCs for simulations carying the pMal-p2x plasmid captured 99% of the cumulative variance in the dataset. Analysis of loadings with an absolute value greater than 90% of the highest loading were recorded and further analysed in Supplementary Figures S4-S6. Several metabolic reactions and subsystems exhibited significant upregulation or downregulation, while some reactions also demonstrated reversed directionality. Among the most notably affected pathways were glycolysis, the tricarboxylic acid (TCA) cycle, and oxidative phosphorylation, all of which were generally upregulated. This pattern suggests an increased cellular demand for ATP, a finding consistent with previous literature [3],[4],[5]. Interestingly, the increased demand for nucleotide building blocks to fuel plasmid replication and expression was observed through the downregulation of nucleotide transport from the cytosol to the periplasm in agreement with experimental findings [3]. In addition, the up- and/or down-regulation of intracellular reactions proved to be sensitive to FBAhop parameter values (*k*_*E*_, *k*_*T*_, *φ*_*P*_). The FBAhop parameters have been introduced to capture different plasmid characteristics such as promoter strength (*k*_*E*_). Therefore, plasmids with different characteristics are expected to lead to varied metabolic outocmes which is reflected in the PCA results.

As intuitively expected, several pathways related to amino acid metabolism were upregulated reflecting the increased demand for protein building blocks. The most significant upregulations were observed in alanine, aspartate as well as branched-chain amino acid (BCAA) metabolism (valine, leucine, and isoleucine). The upregulated pathways are linked to the amino acid composition of the produced recombinant protein product (β-lactamase). However, our analysis highlighted less intuitive metabolic shifts associated with plasmid size and composition. For example, in cells carying the pOri1 plasmid a directionality change of the acetate kinase reaction (EC 2.7.2.1) shifting from net acetate consumption to net acetate production was observed. This is idnicative of overflow-type metabolism under elevated ATP demand and carbon throughput. Cells carying the pOri2 plasmid exhibited altered behavior of the nucleotide salvage pathway, a result consistent with the higher dNTP/NTP demand imposed by the larger plasmid relative to pOri1. Under the increased nucleotide load, our results highlighted a preferential bias toward salvage, potentially because recycling bases and nucleotides is energetically more efficient comapred to de novo synthesis. Furthermore, in cells carying the pOri2 plasmid the directionality of the adenylate kinase (EC 2.7.4.4) catalyzed reaction had shifted, indicating an attempt to compensate the increased resource demands to drive biosynthesis. The absence of an analogous signature in cells carying the pOri1 plasmid suggests that the smaller plasmid does not stress nucleotide homeostasis enough to necessitate salvage reprogramming. Collectively, these findings link plasmid size to nucleotide economy: larger constructs drive salvage-linked adaptations to meet replication-driven nucleotide demand while containing energetic cost, a pattern aligned with known metabolic burden responses in plasmid-bearing bacteria.

Similarly, for cells carrying the pMal-p2x plasmid PCA revealed a clear separation between the metabolic fingerprints of wild type versus plasmid bearing cells (Figure 5). In addition, cells carrying the pMal-p2x plasmid were split in two distinct clusters which upon further inspection revealed distinct metabolic fingerprints between cells that produced high or low amounts of the GI-MBP fusion protein. Cells with elevated GI-MBP translation rates clustered together (Figure 5) due to the increased demand for amino acids which led to amplified flux through glycolysis, the TCA cycle, and oxidative phosphorylation. This translation-driven metabolic displacement is consistent with the observation that minimal expression efficiencies are sufficient to supply mRNA templates, rendering translation the limiting step for GI-MBP yield. The metabolic signature of the high-translation cluster includes higher ATP turnover, intensified precursor drain from central carbon pathways, and pathway-level reallocations that reflect the amino acid cost of sustaining elevated protein production rates. Conversely, models with lower MBP translation rates occupy a neighboring but distinct region of the PC space, indicating a reduced energetic and biosynthetic burden. The consistent separation across three principal components underscores that modulation of translation efficiency creates distinct metabolic phenotypes, even when promoter activity and objective weights on recombinant flux remain modest. Collectively, these results position GI-MBP translation efficiency as the primary driver of phenotypic configuration in the pMal-p2x system, and they highlight how translation-centric control reshapes the host’s energetic economy and pathway utilization to achieve higher recombinant protein productivity (Supplementary Figure S6).

### 3.3 Theoretical Analysis of Process Design Space

The FBAhop algorithm presented herein can accommodate multiple plasmid modalities, ranging from relatively simple constructs expressing a single antibiotic resistance protein (pOri1, pOri2) to more complex plasmids carrying regulatory elements (pMal-p2x). The algorithm introduces parameters related to promoter strength (*k*_*E*_), mRNA translation efficiency (*k*_*T*_) and recombinant protein burden (*φ*_*P*_) thus enabling the investigation of host-plasmid interactions and the optimization of recombinant expression systems. To highlight the insights that can be gained by such an analysis, the Design Space Identification (DSI) algorithm as presented by Sachio et. al [9] was applied for each of the three plasmids considered.

Briefly, FBAhop parameters (*k*_*E*_, *k*_*T*_, *φ*_*P*_) were assigned fixed values (Supplementary Tables S3-S4) and each of the three models was carbon constrained using ccFBA. Maximum theoretical pDNA productivities for pOri1 and pOri2 and maximum theoretical GI-MBP productivity for pMal-p2x were determined by optimizing for the respective flux. DSI was employed to study the tradeoff between increased expression efficiency (*k*_*E*_), growth rate (*v*_*biomass*_) and pDNA or GI-FBP productivity respectively. Expression efficiency (*k*_*E*_) and growth rate (*v*_*biomass*_) were allowed to vary between predefined bounds and 2^15^ combinations were retrieved using a Sobol sequence based random number generator [33]. Simulations were performed for each of the sampled combinations and parameter values that could match a predetermined performance index were recorded. The performance index in the present study was set as a fraction of the maximum theoretical productivity assessed at four distinct levels (30, 50, 70 and 90% of 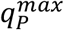).

Cells carrying the pOri1 and pOri2 plasmids were used to simulate pDNA manufacturing. The DSI analysis described above highlighted an inverse correlation between expression efficiency (*k*_*E*_) and pDNA productivity as lower expression efficiency values generally corresponded to higher pDNA fluxes (Figure 6). A lower maximum theoretical productivity was observed for pOri2 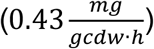 compared to pOri1 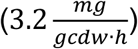, potentially due to the increased plasmid size. Consequently, comparatively higher expression efficiency values (*k*_*E*_) are still compatible with set pDNA productivity levels compared to pOri1. Furthermore, the growth rates at which cells can sustain target pDNA fluxes for pOri2 were lower than those for pOri1, reflecting the amplified metabolic load imposed by the larger plasmid. This observation implies the sensitivity of the pOri2 system to the trade-off between cellular growth and heterologous gene expression, where increased plasmid size and associated metabolic demands intensify resource competition.

**Figure 6.**
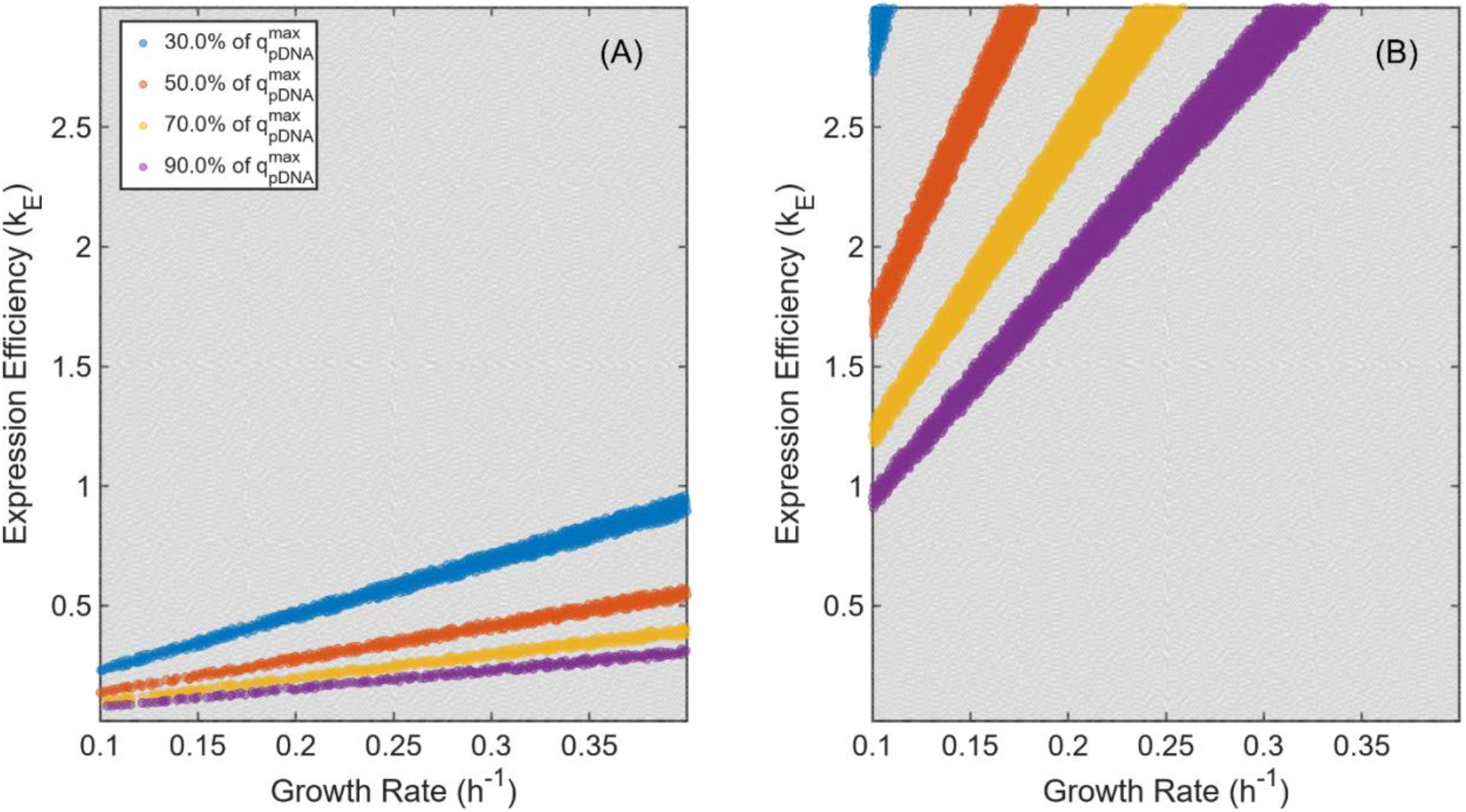
pDNA manufacturing Design Space Identification analysis. Effect of varying expression efficiency (*k*_*E*_) and growth rate (v_biomass_) in order to achieve predefined pDNA productivity targets expressed as a prercentage (30, 50, 70, 90%) of maximum theoretical productivity. Results for cells carrying the: A) pOri1 and B) pOri2 plasmids.

Cells transformed with the pMal-p2x system express the recombinant GI-MBP fusion protein and were used to simulate recombinant protein production. The DSI analysis highlighted that the expression efficiency of the ampicillin resistance gene 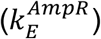 affects the operable design space only at lower values (Figures 7I-L). This suggests that at low 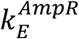 values, tuning of plasmid expression begins to significantly affect cellular growth and, consequently, recombinant protein productivity. Surprisingly, expression efficiencies (*k*_*E*_) for all three expressed genes, including the GI-MBP and lacI genes, did not exert a significant influence on recombinant protein productivity. This counterintuitive result arises because the lowest permissible value set for expression efficiency (0.02 mRNAs per plasmid) allows the production of adequate mRNA templates for protein synthesis under the simulated conditions. A further DSI investigation where translation efficiency (k_T_) was allowed to vary instead of expression efficiency revealed a stronger correlation between 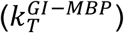 and recombinant protein productivity (Supplementary Figure S9). This aligns with biological expectations, as protein yield is often limited by translation machinery availability rather than transcriptional input alone. Together, these findings reinforce the necessity of considering both transcriptional and translational regulation alongside plasmid-associated metabolic burden when designing and optimizing recombinant expression systems. The trade-offs between plasmid size, expression efficiency, and host growth rates must be carefully balanced to achieve desired productivity targets.

**Figure 7.**
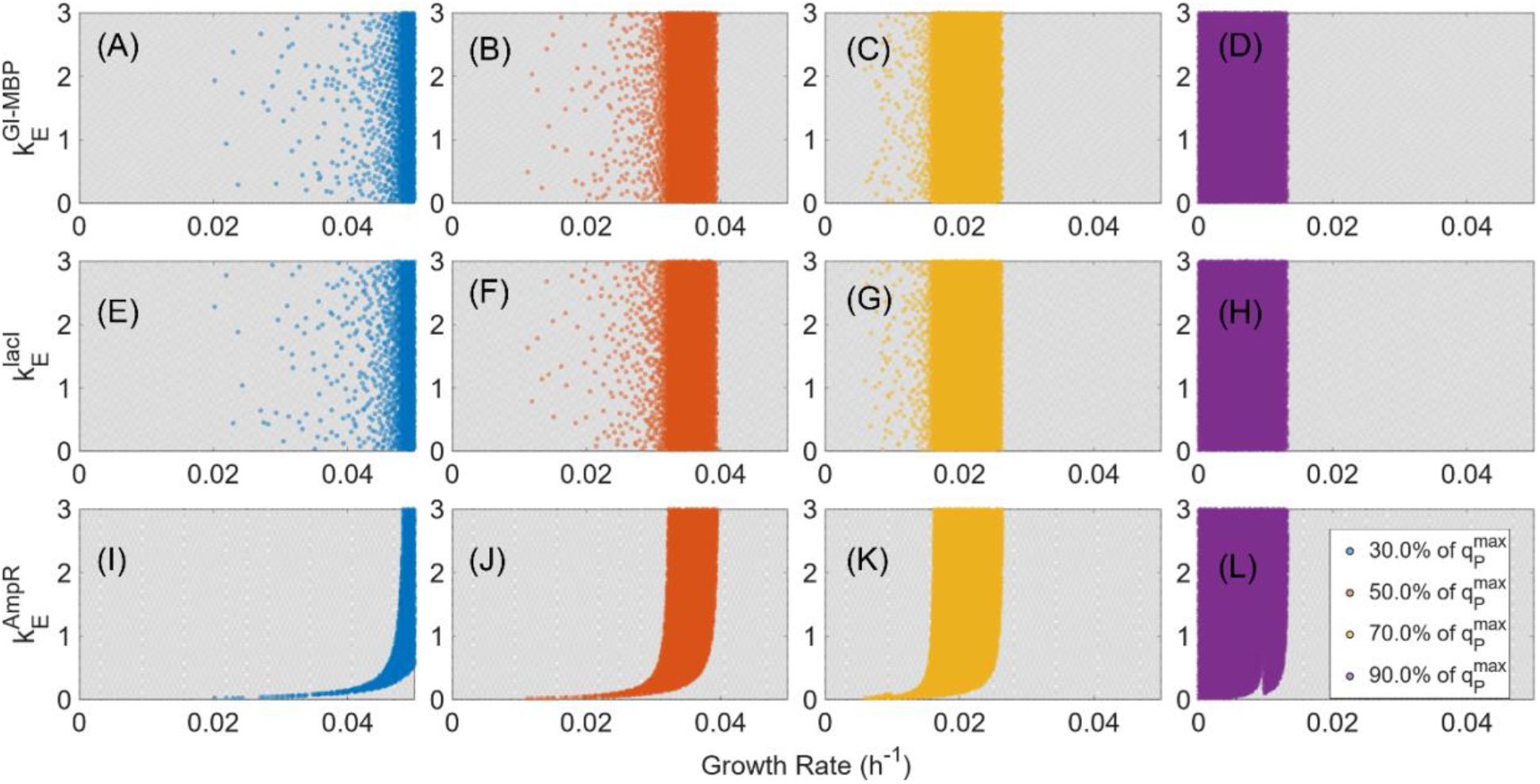
Recombinant protein manufacturing Design Space Identification analysis. Effect of varying expression efficiencies (*k*_*E*_) for each of the three genes on plasmid pMal-p2x and growth rate (v_biomass_) in order to achieve predefined productivity targets expressed as a percentage (30, 50, 70, 90%) of the maximum theoretical productivity. A-D) Impact of MBP-GI expression efficiency 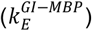 and growth rate (v_biomass_) on recombinant protein productivity; E-H) Impact of lacI expression efficiency 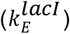 and growth rate (v_biomass_) on recombinant protein productivity and I-L) Impact of AmpR expression efficiency 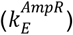 and growth rate (v_biomass_) on recombinant protein productivity.

## Conclusions

A thorough understanding of host-plasmid interactions is necessary for the development of robust, effiecient and high-yield pDNA and recombinant protein manufacturing processes. Several studies have used constraint based modelling approaches in an attempt to capture the metabolic reconfiguration that occurs in host cells in response to the introduction of plasmids [2],[7],[8],[29],[34]. In the present work, we introduced FBAhop, a computational algorithm that quantifies the metabolic impact of plasmid replication and expression by developing tailor made metabolic reactions based on the plasmid’s sequence. By enumerating the metabolic costs from gene sequence, to transcribed mRNA sequence, to final protein product FBAhop effectively captures metabolic shifts that arise from the expression of an heterologous gene, thereby revealing pathway-specific adjustments in organisms transformed with different plasmids.

The impact of three different plasmids of varying size and complexity on *E. coli* metabolism was studied using FBAhop. Our analysis revealed both intuitive and less obvious metabolic shifts that occur is response to variations in plasmid size, heterologous gene expression and translation efficiencies. Detailed model analysis was used to identify key metabolic pathways affected by plasmid integration reflecting the metabolic response to the increased energetic and biosynthetic demands associated with plasmid maintenance and recombinant protein production. Overall, FBAhop provides a robust tool to quantitatively predict the metabolic burden associated with plasmid integration, replication and expression providing valuable insights for the optimization of pDNA and/or recombinant protein production.

## Acknowledgements

Results presented in this work have been produced using the Aristotle University of Thessaloniki (AUTh) High Performance Computing Infrastructure and Resources. This work was funded by the European Research Council under grant agreement No. 101095721 (CoDiBio: Continuous Digitalized Processes for Producing Biopharmaceuticals).

## CRediT authorship contribution statement

**Nikolaos Stratis**: Writing – review & editing, Writing – original draft, Software, Methodology, Investigation, Formal analysis, Data curation, Conceptualization.

**Massimo Morbidelli**: Writing – review & editing, Methodology, Investigation, Conceptualization, Funding acquisition, Supervision, Project administration.

**Alexandros Kiparissides**: Writing – review & editing, Methodology, Investigation, Validation, Supervision, Project administration, Formal analysis, Conceptualization.

