## Supplementary Material 1 for "From plasmid sequence to process design: A computational analysis of metabolism in the context of plasmid DNA manufacturing"

**S1. Comparative performance of *E. coli* GeMs**

^13^C metabolic flux data for intracellular reactions across several central carbon pathways including Glycolysis, Pentose Phosphate Pathway, Entner-Doudoroff Pathway, TCA Cycle, Glyoxylate Shunt, Amphibolic Reactions and Amino Acid Biosynthesis were retrieved from literature [1]. The dataset contained experimentally measured values for intra- and extra-cellular reaction fluxes across different environmental conditions (aerobic and anaerobic) and substrates (Glucose and Xylose). Each GeM was constrained using experimental values for uptake and secretion rates [1], while the specific growth rate was predicted using ccFBA. The total number of reactions across all conditions for which a GeM failed to correctly predict reaction directionality was tracked. For example, if a particular GeM is predicting poistive flux when the experimental flux is negative and vice versa a penalty point is assigned.

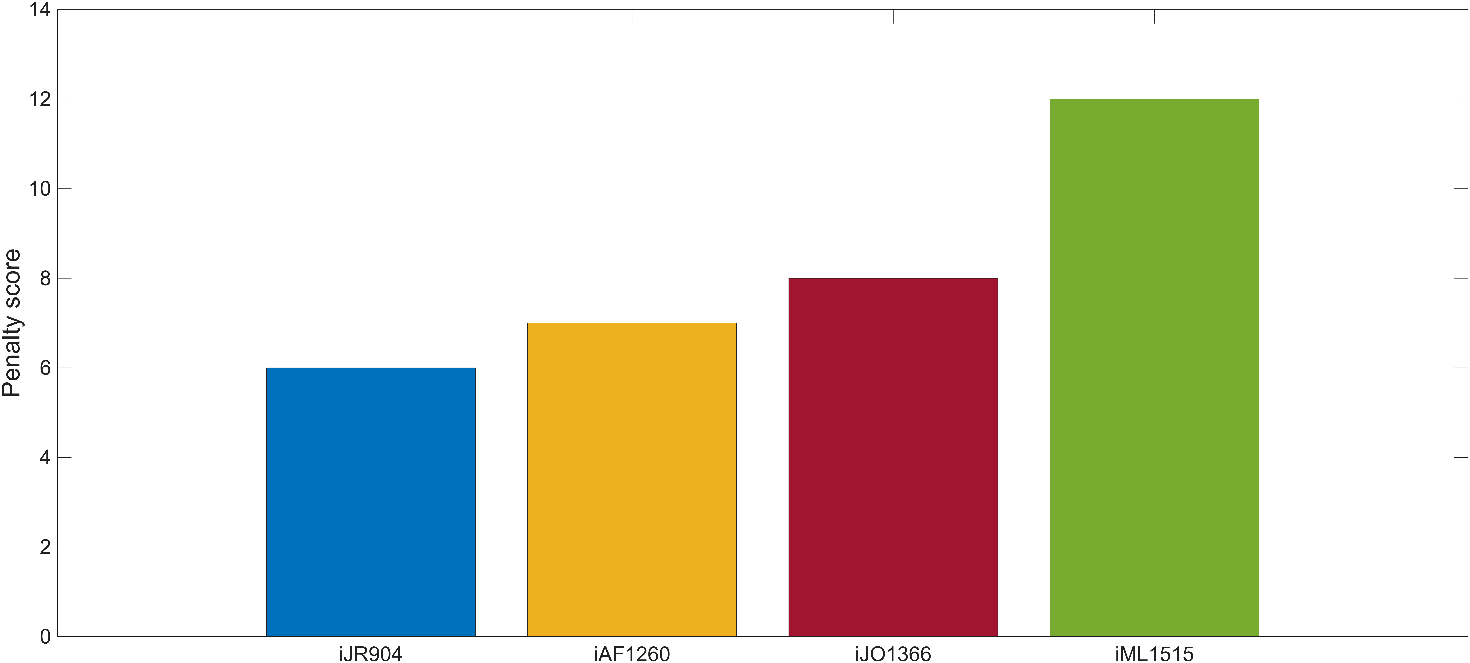

**Supplementary Figure S1:** The total number of reaction directionalities each model failed to predict (Penalty Score)

**S2. Global Parametric Analysis of FBAhop parameters**

For the pMal-p2x plasmid, $k_{E}$was set between 0.02 and 3 mRNA molecules per gene ($\frac{mRNA}{gene}$) [2], $k_{T}$was set between 10 and 100 protein molecules per mRNA molecule ($\frac{P}{mRNA}$) [3] and $\varphi_{P}$ was set between 0 and 12% [4] total protein content ($\frac{g_{P}}{g_{P,Total}}$) . The parameters were then allowed to vary simultaneously within the above defined ranges. A total of 2^20^ points were sampled from the resulting multi-dimensional parameter space using a Sobol sequence based random number generator [5]. The pMal-p2x model was simulated once for each set of sampled parameter values and parameter sets that accurately predicted the experimentally reported growth rate (Table 3) were recorded. Their distributions are shown below

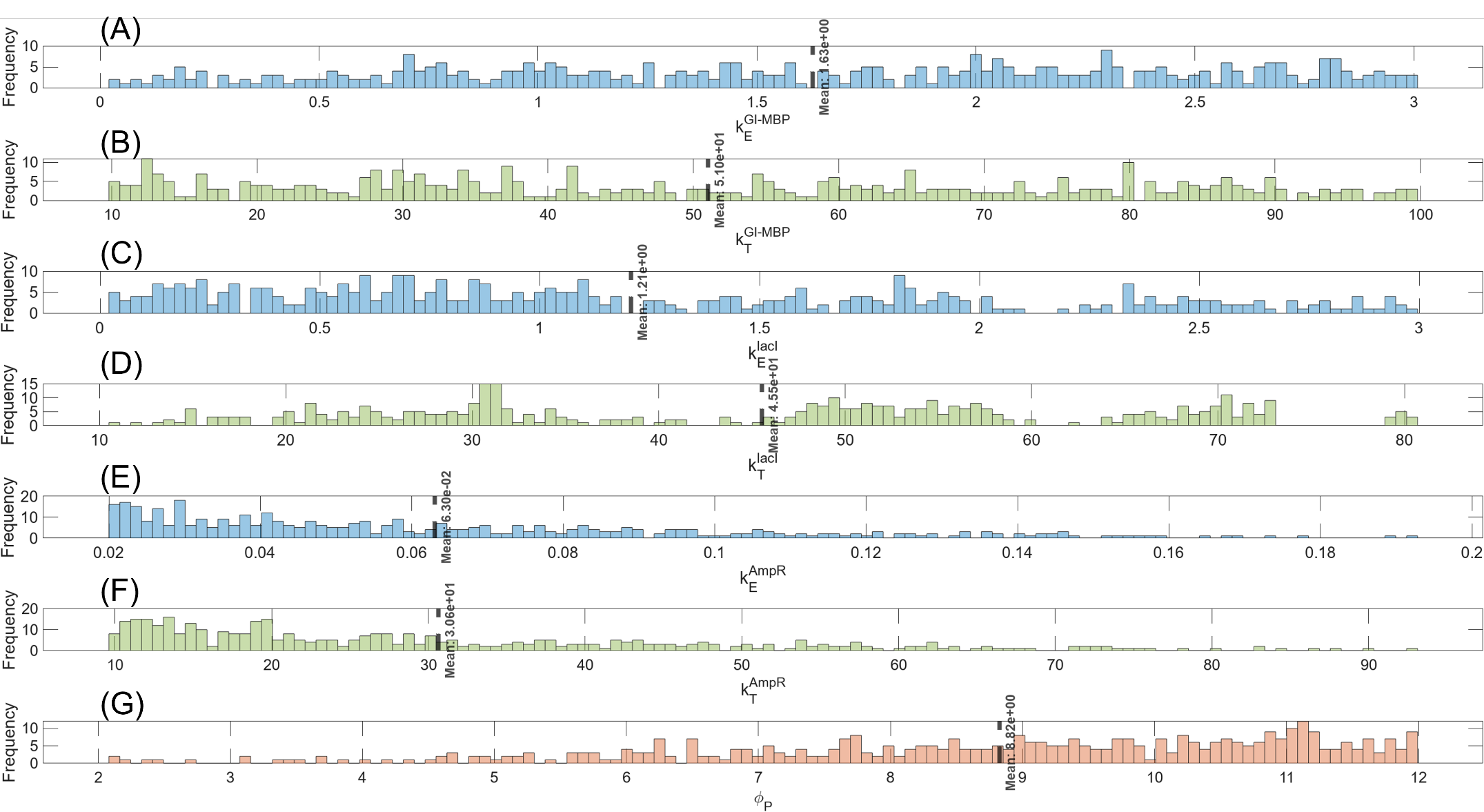

**Supplementary Figure S2:** Global parametric analysis of FBAhop parameters for plasmid pMal-p2x. Distributions of randomly sampled values that can accurately predict experimentally reported growth rates and GI-MBP flux: A, C, E) expression efficiency (kE Eq.7) for the β-lactamase, laqI and GI-MBP encoding genes respectively; B, D, F) translation efficiency (kT Eq.8) for the β-lactamase, laqI and GI-MBP mRNAs respectively and G) fraction of total protein corresponding to the antibiotic resistance protein (φ_P_).

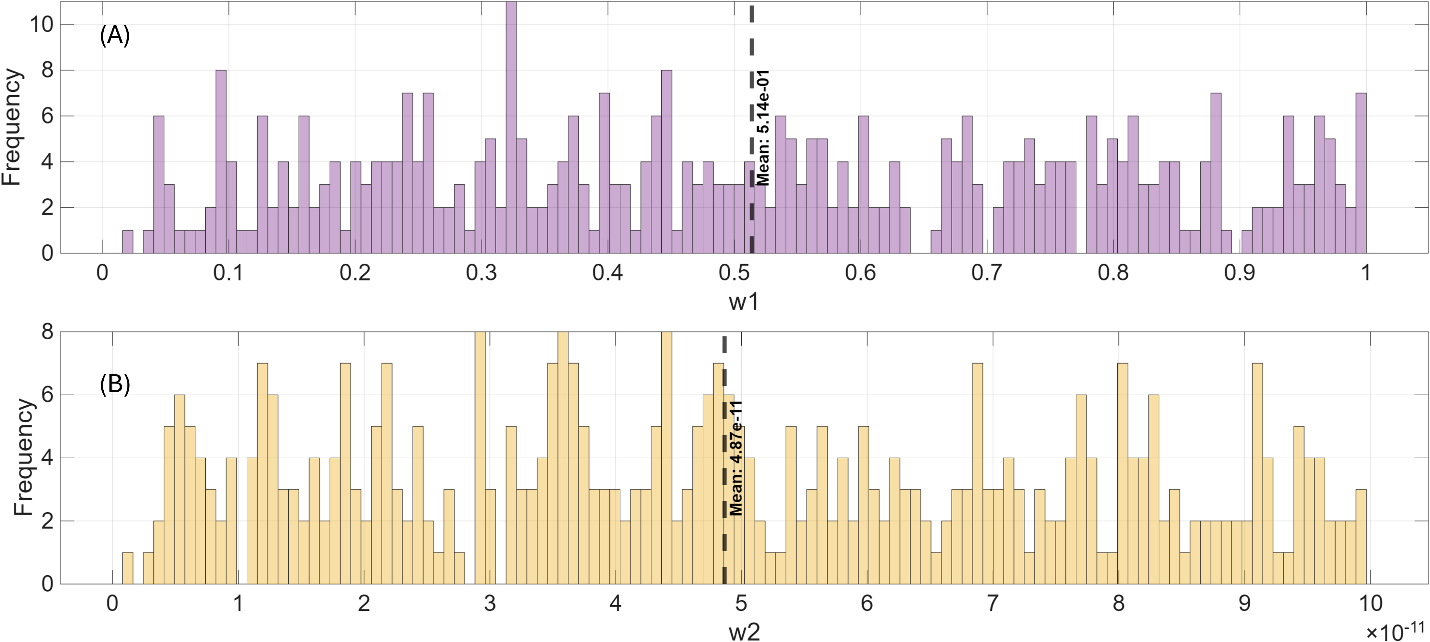

**Supplementary Figure S3:** Global parametric analysis of FBAhop parameters for plasmid pMal-p2x. Distributions of randomly sampled values that can accurately predict experimentally reported growth rates: A) Biomass weight factor (w_1_) in the objective function formulation (Eq. 15) and B) Recombinant fusion protein weight factor (w_2_) in the objective function formulation (Eq. 15).

**S3. Principal Component Analysis**

To further understand the impact and functional role of the parameters introduced by FBAhop a series of *in silico* experiments was performed. Specifically, a 3-level Box Behnken Design was employed to generate sets of parameter values for expression (*k_E_*), translation (*k_T_*) and recombinant protein weight fraction (*φ_P_*) parameters. The levels for the experimental design were based on the sampled distribution means ± one standard deviation (Figures 2, 3 and Supplemental Figures S2, S3). Each of the parameter sets defined by the experimental design (Supplemental Tables S1, S2) were simulated separately.

**Supplementary Table S1:** Box Behnken Design of FBAhop parameters for the simulation of E. coli cells carrying the pOri1 and pOri2 plasmids

| **Model** | **pOri1** | | | **pOri2** | | |
| --- | --- | --- | --- | --- | --- | --- |
|  | ***k_E_*** | ***φ_P_*** | ***k_T_*** | ***k_E_*** | ***φ_P_*** | ***k_T_*** |
| A | 0.08 | 10 | 37 | 0.1 | 10 | 37 |
| B | 0.08 | 5.5 | 37 | 0.1 | 5 | 37 |
| C | 0.08 | 7.75 | 12 | 0.1 | 7.5 | 12 |
| D | 0.08 | 7.75 | 62 | 0.1 | 7.5 | 62 |
| E | 0.44 | 10 | 12 | 0.55 | 10 | 12 |
| F | 0.44 | 10 | 62 | 0.55 | 10 | 62 |
| G | 0.44 | 5.5 | 12 | 0.55 | 5 | 12 |
| H | 0.44 | 5.5 | 62 | 0.55 | 5 | 62 |
| I | 0.44 | 7.75 | 37 | 0.55 | 7.5 | 37 |
| J | 0.44 | 7.75 | 37 | 0.55 | 7.5 | 37 |
| K | 0.44 | 7.75 | 37 | 0.55 | 7.5 | 37 |
| L | 0.8 | 10 | 37 | 1 | 10 | 37 |
| M | 0.8 | 5.5 | 37 | 1 | 5 | 37 |
| N | 0.8 | 7.75 | 12 | 1 | 7.5 | 12 |
| O | 0.8 | 7.75 | 62 | 1 | 7.5 | 62 |

**Supplementary Table S2**: Box Behnken Design of FBAhop parameters for the simulation of E. coli cells carrying the pMal-p2x plasmid

| **Gene** | **lacI** | | **GI-MBP** | | **AmpR** | | |
| --- | --- | --- | --- | --- | --- | --- | --- |
| **Model** | ***k_E_*** | ***k_T_*** | ***k_E_*** | ***k_T_*** | ***k_E_*** | ***k_T_*** | ***φ_P_*** |
| 1 | 1.6275 | 50.9983 | 1.207 | 30.9582 | 0.0324 | 15.6007 | 8.8237 |
| 2 | 1.6275 | 50.9983 | 1.207 | 30.9582 | 0.0324 | 45.6713 | 8.8237 |
| 3 | 1.6275 | 50.9983 | 1.207 | 30.9582 | 0.0936 | 15.6007 | 8.8237 |
| 4 | 1.6275 | 50.9983 | 1.207 | 30.9582 | 0.0936 | 45.6713 | 8.8237 |
| 5 | 1.6275 | 50.9983 | 1.207 | 60.1064 | 0.0324 | 15.6007 | 8.8237 |
| 6 | 1.6275 | 50.9983 | 1.207 | 60.1064 | 0.0324 | 45.6713 | 8.8237 |
| 7 | 1.6275 | 50.9983 | 1.207 | 60.1064 | 0.0936 | 15.6007 | 8.8237 |
| 8 | 1.6275 | 50.9983 | 1.207 | 60.1064 | 0.0936 | 45.6713 | 8.8237 |
| 9 | 0.9692 | 50.9983 | 1.207 | 45.5323 | 0.063 | 15.6007 | 7.0199 |
| 10 | 0.9692 | 50.9983 | 1.207 | 45.5323 | 0.063 | 15.6007 | 10.6275 |
| 11 | 0.9692 | 50.9983 | 1.207 | 45.5323 | 0.063 | 45.6713 | 7.0199 |
| 12 | 0.9692 | 50.9983 | 1.207 | 45.5323 | 0.063 | 45.6713 | 10.6275 |
| 13 | 2.2858 | 50.9983 | 1.207 | 45.5323 | 0.063 | 15.6007 | 7.0199 |
| 14 | 2.2858 | 50.9983 | 1.207 | 45.5323 | 0.063 | 15.6007 | 10.6275 |
| 15 | 2.2858 | 50.9983 | 1.207 | 45.5323 | 0.063 | 45.6713 | 7.0199 |
| 16 | 2.2858 | 50.9983 | 1.207 | 45.5323 | 0.063 | 45.6713 | 10.6275 |
| 17 | 1.6275 | 30.0696 | 1.207 | 45.5323 | 0.0324 | 30.636 | 7.0199 |
| 18 | 1.6275 | 30.0696 | 1.207 | 45.5323 | 0.0324 | 30.636 | 10.6275 |
| 19 | 1.6275 | 30.0696 | 1.207 | 45.5323 | 0.0936 | 30.636 | 7.0199 |
| 20 | 1.6275 | 30.0696 | 1.207 | 45.5323 | 0.0936 | 30.636 | 10.6275 |
| 21 | 1.6275 | 71.927 | 1.207 | 45.5323 | 0.0324 | 30.636 | 7.0199 |
| 22 | 1.6275 | 71.927 | 1.207 | 45.5323 | 0.0324 | 30.636 | 10.6275 |
| 23 | 1.6275 | 71.927 | 1.207 | 45.5323 | 0.0936 | 30.636 | 7.0199 |
| 24 | 1.6275 | 71.927 | 1.207 | 45.5323 | 0.0936 | 30.636 | 10.6275 |
| 25 | 0.9692 | 30.0696 | 1.207 | 30.9582 | 0.063 | 30.636 | 8.8237 |
| 26 | 0.9692 | 30.0696 | 1.207 | 60.1064 | 0.063 | 30.636 | 8.8237 |
| 27 | 0.9692 | 71.927 | 1.207 | 30.9582 | 0.063 | 30.636 | 8.8237 |
| 28 | 0.9692 | 71.927 | 1.207 | 60.1064 | 0.063 | 30.636 | 8.8237 |
| 29 | 2.2858 | 30.0696 | 1.207 | 30.9582 | 0.063 | 30.636 | 8.8237 |
| 30 | 2.2858 | 30.0696 | 1.207 | 60.1064 | 0.063 | 30.636 | 8.8237 |
| 31 | 2.2858 | 71.927 | 1.207 | 30.9582 | 0.063 | 30.636 | 8.8237 |
| 32 | 2.2858 | 71.927 | 1.207 | 60.1064 | 0.063 | 30.636 | 8.8237 |
| 33 | 1.6275 | 50.9983 | 0.5481 | 30.9582 | 0.063 | 30.636 | 7.0199 |
| 34 | 1.6275 | 50.9983 | 0.5481 | 30.9582 | 0.063 | 30.636 | 10.6275 |
| 35 | 1.6275 | 50.9983 | 0.5481 | 60.1064 | 0.063 | 30.636 | 7.0199 |
| 36 | 1.6275 | 50.9983 | 0.5481 | 60.1064 | 0.063 | 30.636 | 10.6275 |
| 37 | 1.6275 | 50.9983 | 1.8659 | 30.9582 | 0.063 | 30.636 | 7.0199 |
| 38 | 1.6275 | 50.9983 | 1.8659 | 30.9582 | 0.063 | 30.636 | 10.6275 |
| 39 | 1.6275 | 50.9983 | 1.8659 | 60.1064 | 0.063 | 30.636 | 7.0199 |
| 40 | 1.6275 | 50.9983 | 1.8659 | 60.1064 | 0.063 | 30.636 | 10.6275 |
| 41 | 0.9692 | 50.9983 | 0.5481 | 45.5323 | 0.0324 | 30.636 | 8.8237 |
| 42 | 0.9692 | 50.9983 | 0.5481 | 45.5323 | 0.0936 | 30.636 | 8.8237 |
| 43 | 0.9692 | 50.9983 | 1.8659 | 45.5323 | 0.0324 | 30.636 | 8.8237 |
| 44 | 0.9692 | 50.9983 | 1.8659 | 45.5323 | 0.0936 | 30.636 | 8.8237 |
| 45 | 2.2858 | 50.9983 | 0.5481 | 45.5323 | 0.0324 | 30.636 | 8.8237 |
| 46 | 2.2858 | 50.9983 | 0.5481 | 45.5323 | 0.0936 | 30.636 | 8.8237 |
| 47 | 2.2858 | 50.9983 | 1.8659 | 45.5323 | 0.0324 | 30.636 | 8.8237 |
| 48 | 2.2858 | 50.9983 | 1.8659 | 45.5323 | 0.0936 | 30.636 | 8.8237 |
| 49 | 1.6275 | 30.0696 | 0.5481 | 45.5323 | 0.063 | 15.6007 | 8.8237 |
| 50 | 1.6275 | 30.0696 | 0.5481 | 45.5323 | 0.063 | 45.6713 | 8.8237 |
| 51 | 1.6275 | 30.0696 | 1.8659 | 45.5323 | 0.063 | 15.6007 | 8.8237 |
| 52 | 1.6275 | 30.0696 | 1.8659 | 45.5323 | 0.063 | 45.6713 | 8.8237 |
| 53 | 1.6275 | 71.927 | 0.5481 | 45.5323 | 0.063 | 15.6007 | 8.8237 |
| 54 | 1.6275 | 71.927 | 0.5481 | 45.5323 | 0.063 | 45.6713 | 8.8237 |
| 55 | 1.6275 | 71.927 | 1.8659 | 45.5323 | 0.063 | 15.6007 | 8.8237 |
| 56 | 1.6275 | 71.927 | 1.8659 | 45.5323 | 0.063 | 45.6713 | 8.8237 |
| 57 | 1.6275 | 50.9983 | 1.207 | 45.5323 | 0.063 | 30.636 | 8.8237 |
| 58 | 1.6275 | 50.9983 | 1.207 | 45.5323 | 0.063 | 30.636 | 8.8237 |
| 59 | 1.6275 | 50.9983 | 1.207 | 45.5323 | 0.063 | 30.636 | 8.8237 |
| 60 | 1.6275 | 50.9983 | 1.207 | 45.5323 | 0.063 | 30.636 | 8.8237 |
| 61 | 1.6275 | 50.9983 | 1.207 | 45.5323 | 0.063 | 30.636 | 8.8237 |
| 62 | 1.6275 | 50.9983 | 1.207 | 45.5323 | 0.063 | 30.636 | 8.8237 |

Each of the parameter sets defined by the experimental design (Supplemental Tables S1, S2) were simulated separately. The resulting models were carbon constrained [6] and 10,000 solution vectors were retrieved from each model’s solution space using a Markov-Chain Monte Carlo algorithm. This resulted in 15 unique models simulating *E. coli* cells carrying the pOri1 plasmid, 15 unique models for cells carrying the pOri2 plasmid and 62 unique models for cells carrying

the pMal-p2x plasmid.

Principal Component Analysis (PCA) was employed to systematically assess the metabolic reconfiguration of host cells following plasmid transformation and recombinant product expression following the methodology presented by [7]. Briefly, the sampled solution vectors from the models generated by the experimental design were allocated in data pools according to the carried plasmid. Each data pool was augmented by the addition of solution vectors from wild type *E. coli* carrying no plasmid. PCA was applied individually to each data pool and loadings with a value higher than 90% of the highest loading were recorded for further analysis. A heatmap of the reactions with substantially different behavior compared to the wild type for each of the simulations defined in Supplementary Tables 1 and 2 is presented in Supplementary Figures 4-6. Red color shows down regulation, green upregulation and black reversed directionality in respect to the reactions mean flux distribution simulated in the Wild Type (BL21).

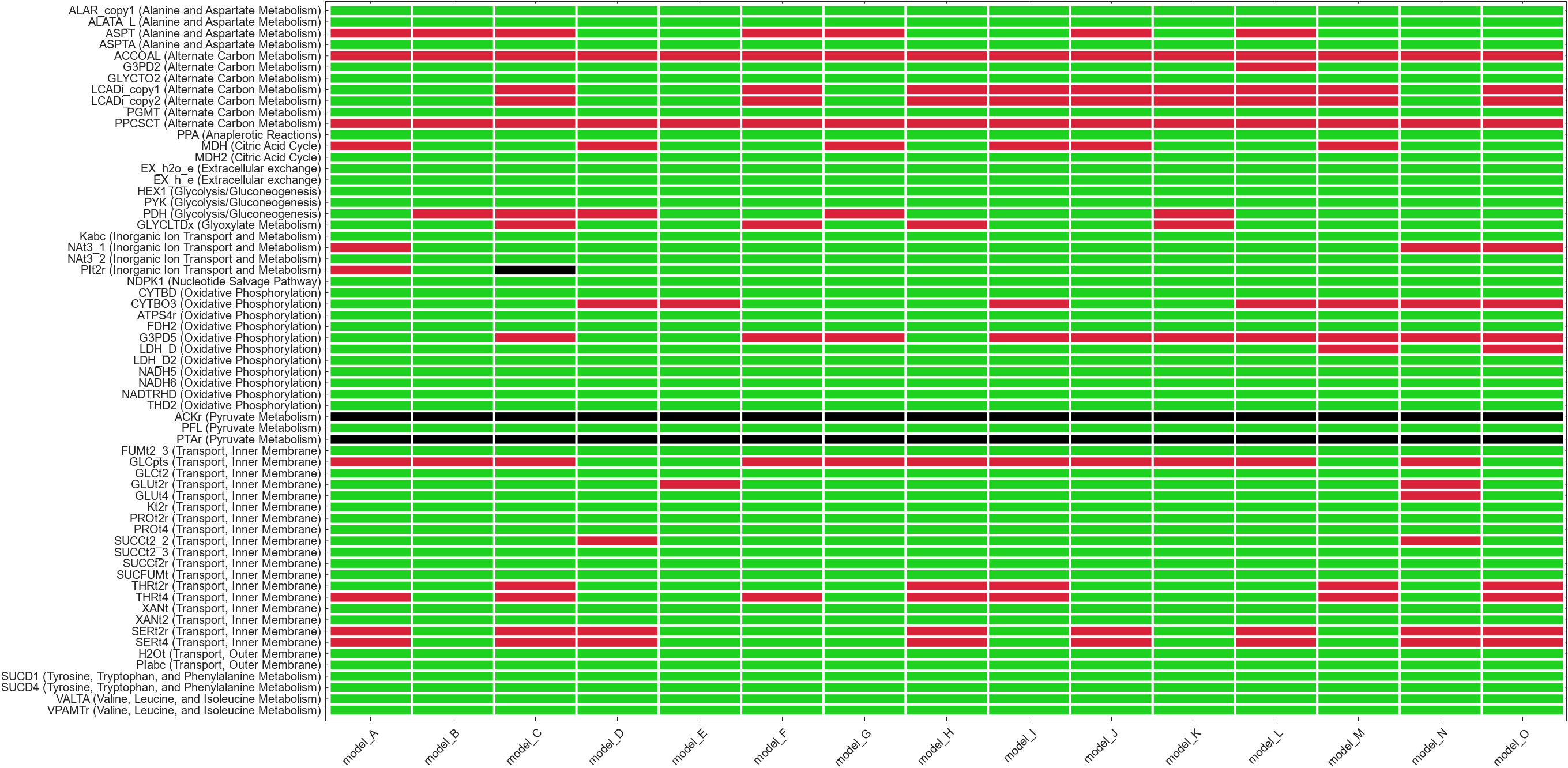

**Supplementary Figure S4:** Metabolic fingerprint of plasmid bearing E. coli cells. Up- or down-regulation of reactions with high loadings during principal component analysis of sampled solution vectors retrieved from both wild type and pOri1 plasmid bearing simulations. Red colour represents downregulation, green upregulation and black indicates a reversed directionallity with respect to wild type behavior.

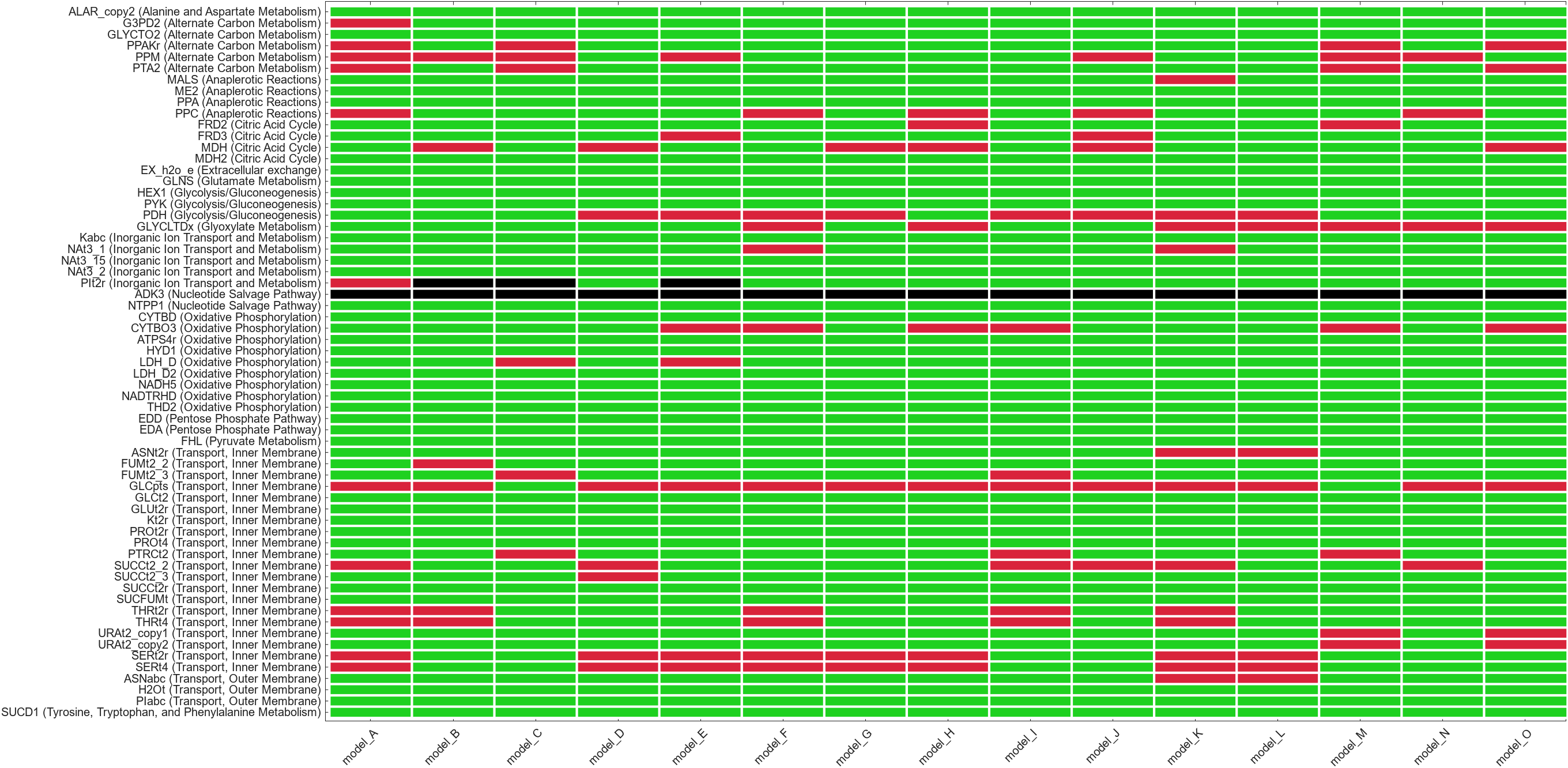

**Figure S5** Metabolic fingerprint of plasmid bearing E. coli cells. Up- or down-regulation of reactions with high loadings during principal component analysis of sampled solution vectors retrieved from both wild type and pOri2 plasmid bearing simulations. Red colour represents downregulation, green upregulation and black indicates a reversed directionallity with respect to wild type behavior.

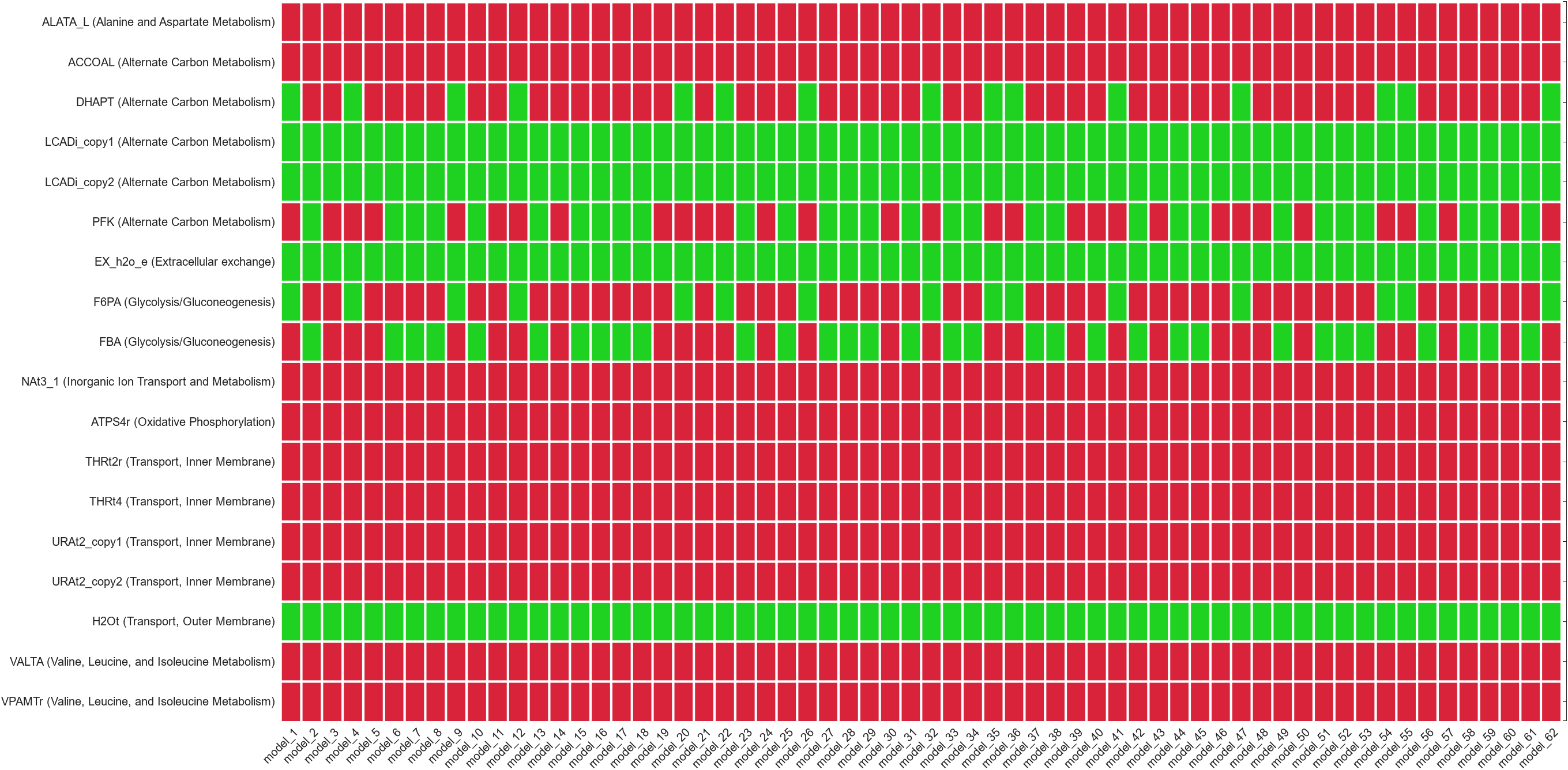

**Figure S6** Metabolic fingerprint of plasmid bearing E. coli cells. Up- or down-regulation of reactions with high loadings during principal component analysis of sampled solution vectors retrieved from both wild type and pMal-p2x plasmid bearing simulations. Red colour represents downregulation, green upregulation and black indicates a reversed directionallity with respect to wild type behavior

**S4. Calculation of maximum theoretical pDNA and/or recombinant protein productivities**

For cells carrying the pOri1 and pOri2 plasmids the impact of varying levels of expression ($k_{E}$) and translation ($k_{T}$) efficiencies at predefined levels of antibiotic resistance protein productivity ($\varphi_{P}$) were investigated. Figures S7 (for pOri1) and S8 (for pOri2) highlight that lower $k_{E}$ values (moles of mRNA per mole of plasmid) lead to higher pDNA productivity for the same translation efficiency ($k_{T}$) and recombinant protein weight fraction levels ($\varphi_{P}$). This means that when a promoter is weak (low mRNA per plasmid levels), cells must generate greater quantities of pDNA to produce a sufficient amount of resistance protein for survival. Notably, as $\varphi_{P}$ increases, the infeasibility area (white regions), representing parameter combinations where the ccFBA problem fails to solve, expands. In these regions, the cellular resource allocation becomes unsustainable, meaning that the metabolic and energetic demands imposed by plasmid maintenance and recombinant protein expression exceed the cell’s capacity, preventing viable growth.

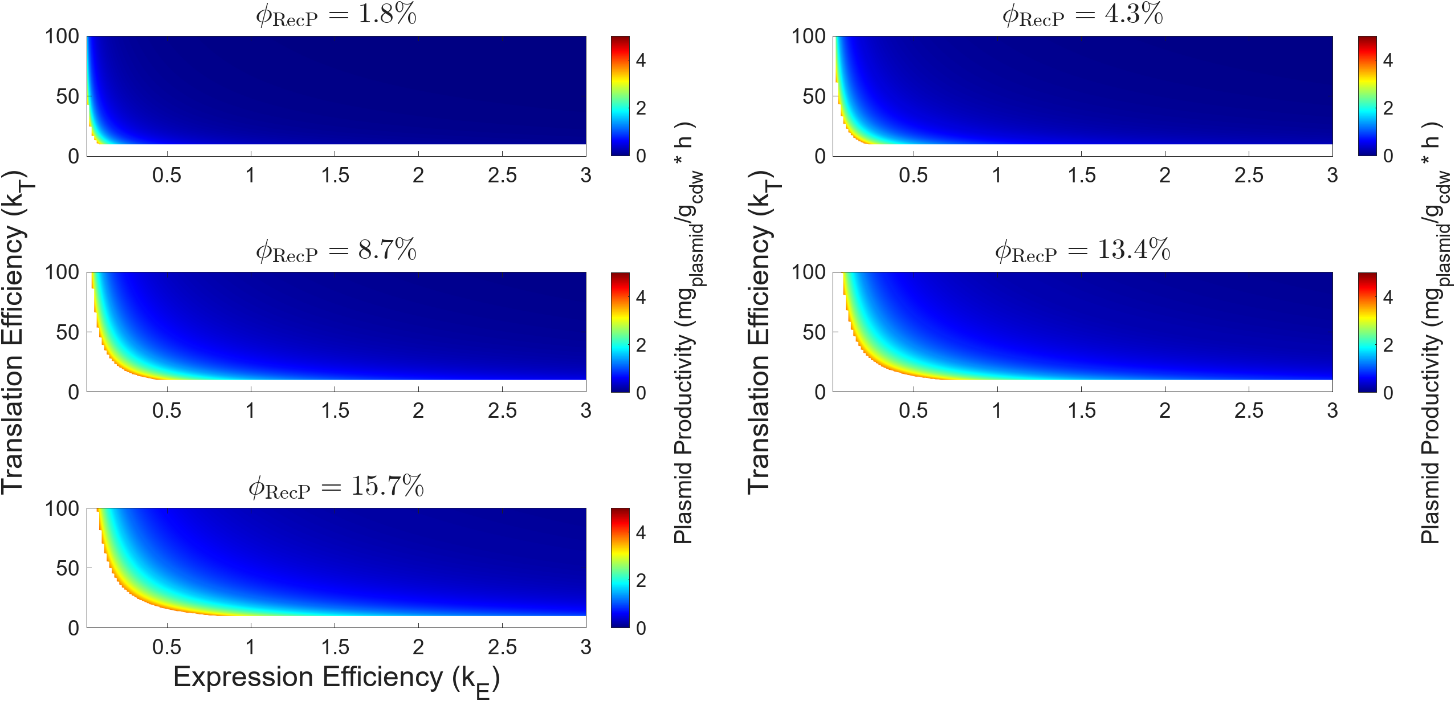
**Supplementary Figure S7:** Effect of expression ($k_{E}$) and translation ($k_{T}$) efficiency on pDNA productivity for cells carrying the pOri1 plasmid. Growth rate is fixed at (v_biomass_ = 0.30), $\varphi_{P}$ is fixed at predefined levels.

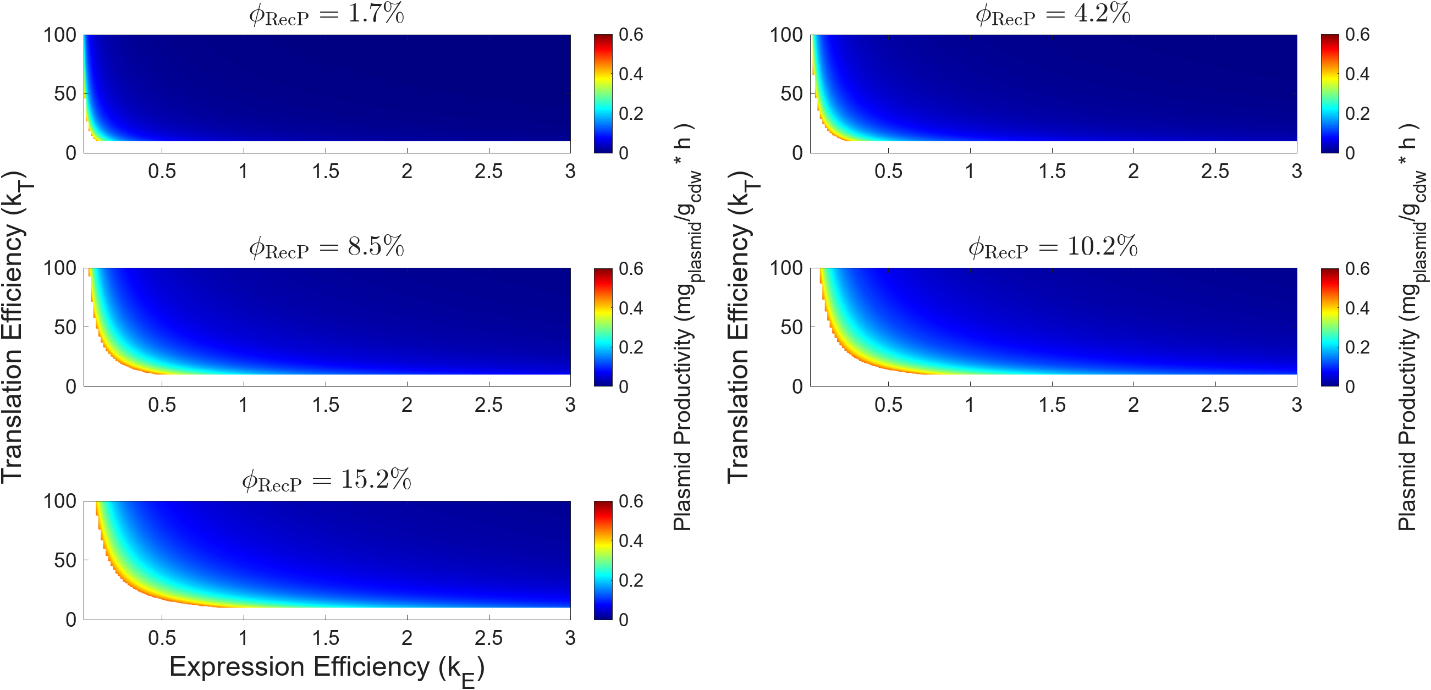

**Supplementary Figure S8:** Effect of expression ($k_{E}$) and translation ($k_{T}$) efficiency on pDNA productivity for cells carrying the pOri2 plasmid. Growth rate is fixed at (v_biomass_ = 0.20), $\varphi_{P}$ is fixed at predefined levels.

Based on the analysis presented above the following values were selected in order to compute the maximum theoretical pDNA productivity:

**Supplementary Table S3:** FBAhop parameter values used to calculate maximum theoretical pDNA productivity

|  | **pOri1** | | | **pOri2** | | |
| --- | --- | --- | --- | --- | --- | --- |
|  | ***k_E_*** | ***k_T_*** | ***φ_P_*** | ***k_E_*** | ***k_T_*** | ***φ_P_*** |
| Value | 0.02 | 37 | 8.7 | 0.02 | 37 | 8.5 |

For cells carrying the pMal-p2x plasmid the following values were selected in order to compute the maximum theoretical recombinant protein productivity:

**Supplementary Table S4:** FBAhop parameter values used to calculate maximum theoretical recombinant protein productivity

|  | **lacI** | | **GI-MBP** | | **AmpR** | | |
| --- | --- | --- | --- | --- | --- | --- | --- |
|  | ***k_E_*** | ***k_T_*** | ***k_E_*** | ***k_T_*** | ***k_E_*** | ***k_T_*** | ***φ_P_*** |
| Value | 1.2 | 45 | 1.6 | 51 | 0.06 | 36 | 8.8 |

**S4. Theoretical Analysis of Process Design Space**

Design space identification for cells carrying the pMal-p2x plasmid where translation efficiencies (*k_T_*) and growthrate (*v_biomass_*) were allowed to vary between predefined bounds.

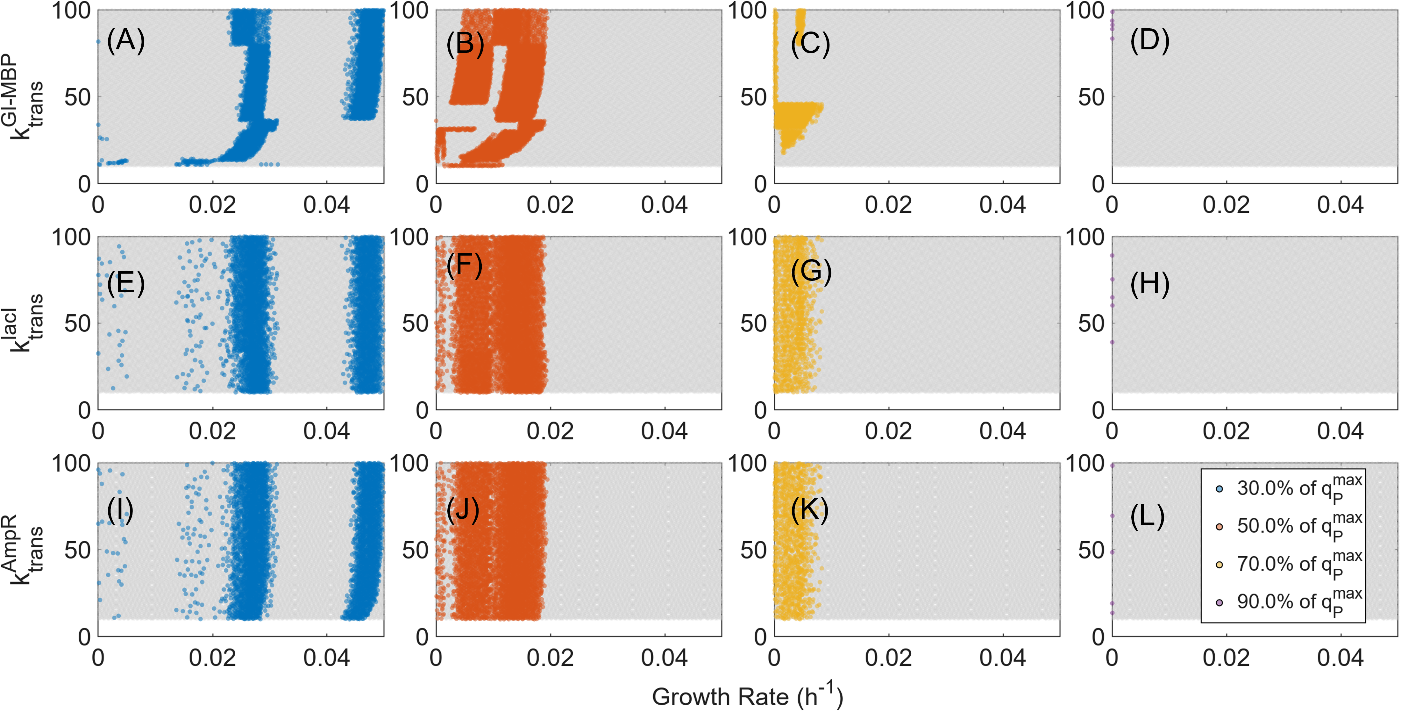

**Supplementary Figure S9** Recombinant protein manufacturing Design Space Identification analysis. Effect of varying translation efficiencies (k_T_) for each of the three genes on plasmid pMal-p2x and growthrate (v_biomass_) in order to achieve predefined productivity targets expressed as a percentage (30, 50, 70, 90%) of the maximum theoretical productivity. A-D) Impact of MBP-GI expression efficiency ($k_{E}^{MBP-GI}$) and growthrate (v_biomass_) on recombinant protein productivity; E-H) Impact of lacI expression efficiency ($k_{E}^{lacI}$) and growthrate (v_biomass_) on recombinant protein productivity and I-L) Impact of AmpR expression efficiency ($k_{E}^{AmpR}$) and growthrate (v_biomass_) on recombinant protein productivity.
